# Rapid estimation of receptive fields across stages of the early visual system

**DOI:** 10.64898/2026.09.16.752021

**Authors:** Marc Büttner, Matej Znidaric, Roland Diggelmann, Federica B. Rosselli, Annalisa Bucci, Andreas Hierlemann, Felix Franke

## Abstract

Estimating receptive fields in large neural populations is often limited by experimental time: stimuli with theoretically favorable statistics for receptive-field estimation may, in practice, evoke too few informative spikes in neurons with high specificity for structured features, requiring impracticably long recordings. Therefore, we asked whether a structured stimulus could broadly engage neural populations across stages of the early visual system while permitting a tractable and interpretable reverse-correlation analysis. To address this question, we developed “reverse correlation against stimulus elements (RCASE)”, an analysis framework for stimuli that can be represented frame by frame through the presence or absence of specific stimulus elements. We then designed such a stimulus consisting of random moving objects (RMO) and compared RMO/RCASE with conventional spike-triggered averaging using a dense binary white noise (WN/STA) stimulus across different stages of the early visual system of the mouse, namely the retina, the nucleus of the optic tract, the superior colliculus and the primary visual cortex, and in the primate retina. To distinguish the contributions of spatial and temporal scales, motion, stimulus structure, and analysis choice, we additionally compared against WN/STA with different square sizes and frame rates, spike-triggered covariance analysis, as well as random static objects and spatially correlated cloud stimuli. RMO evoked stronger and more reliable responses than WN in the mouse retina and across the investigated mouse visual areas. RMO/RCASE yielded discernible receptive fields for a substantially larger proportion of neurons and in a fraction of the recording time. Although the stimulus objects featured localized moving contrast edges, this advantage was not restricted to direction-selective cell populations. In the primate retina, in contrast, WN/STA approached the performance of RMO/RCASE. RMO/RCASE also supported simultaneous estimation of stimulus direction, speed, and color tuning. Our approach enables rapid and interpretable functional characterization of large populations of visual neurons, enabling to simultaneously recover receptive fields and feature tuning, whenever a low-dimensional feature space can be specified beforehand.

## Introduction

Visual systems comprise parallel pathways and cell types that are specialized for different aspects of a visual scene (Nassi & Callaway, 2009). This functional diversity is already prominent in the retina, where retinal ganglion cell (RGC) types differ in their sensitivity to contrast, spatial scale, temporal dynamics, and motion direction (Dhande et al., 2013; Farrow et al., 2013; Krieger et al., 2017; Sabbah et al., 2017; A. Y. M. Wang et al., 2023), and continues in downstream components of the visual system. Experimentally, it is often favorable to estimate the receptive-field locations of as many recorded neurons as possible in the shortest possible time. To achieve this feat, the employed visual stimulus must engage many neurons simultaneously, sample the relevant stimulus dimensions across the visual field, and provide sufficient data for interpretable receptive-field estimates within the limited duration of an experiment.

A receptive field (RF) is necessarily defined relative to a stimulus representation. Pixel-based analyses describe sensitivity to local luminance or contrast, whereas feature-based analyses can describe sensitivity to properties, such as spatial location, motion direction, or speed. The stimulus representation, therefore, determines both what can be estimated and how efficiently it can be estimated. Stimuli, such as dense white noise (WN), broadly sample the stimulus space without requiring specific stimulus features to be specified in advance but may evoke few informative responses from neurons that specifically react to such structured features. Conversely, a more narrowly defined stimulus representation can concentrate sampling on hypothesized features, at the cost that responses to features outside that representation remain hidden. Dense WN remains an important reference for RF estimation, because its controlled statistics support parallel mapping of RFs across neural populations under well-defined assumptions using spike-triggered averaging (STA) (Baccus & Meister, 2002; Chichilnisky, 2001; Park & Pillow, 2011; Rieke, 2001; D. Ringach & Shapley, 2004; Schwartz et al., 2006). These WN/STA RFs describe the linear component of the neural response as a function of pixel contrast. Structured stimuli instead concentrate stimulus sampling or analysis inside a hypothesized region of the stimulus space. Examples include reverse correlation against parameterized bars (Hubel & Wiesel, 1959, 1962), estimation from naturalistic stimuli (Baudot et al., 2013; D. L. Ringach et al., 2002; T. Sharpee et al., 2004; T. O. Sharpee et al., 2006; Theunissen et al., 2001), and analyses using nonlinear feature representations (T. Sharpee et al., 2004; Shi et al., 2019; Skriabine et al., 2026; Wu et al., 2006). Feature-space characterizations do not need to be readily interpretable as functions of pixel contrast and may not even include a spatial dimension. Characterizations of neural responses as functions of selected feature parameters are commonly referred to as tuning functions. A direction-tuning curve, for example, describes response magnitude as a function of stimulus direction. Here we ask a practical question: Can a structured, parameterized stimulus, combined with a corresponding feature-based analysis, yield interpretable RF estimates for more neurons than conventional WN/STA within a fixed experimental time? To address this question, we developed a method, termed “reverse correlation against stimulus elements (RCASE)”, in which each stimulus frame is encoded through the presence or absence of stimulus elements defined by prespecified parameters, such as the position, direction, color, and speed of moving objects. RCASE then estimates response kernels linearly in this stimulus-element space.

Since the mapping from display pixels to stimulus elements is nonlinear and non-invertible, the resulting neural response kernels cannot generally be interpreted as linear pixel-space RFs. RCASE instead characterizes neuronal tuning to a prespecified, low-dimensional set of stimulus features, and the recovered kernels are directly interpretable in terms of those features.

We tested the developed framework using random moving objects (RMO), a stimulus in which multiple white squares appear at random locations and move along straight trajectories in random directions. RMO thereby combines broad spatial sampling with localized moving contrast edges. As object position is encoded in the RCASE element representation, the resulting RCASE kernel contains a spatial component expressed in display coordinates alongside tuning to other object parameters. RMO/RCASE can, therefore, recover a spatial RF estimate together with tuning functions for parameters, such as direction, speed, and color.

To test RMO/RCASE in neural populations with widely differing response properties, we measured neural responses to WN, RMO, and a battery of other stimuli in the mouse visual system, i.e., the retina, the nucleus of the optic tract (NOT), the superior colliculus (SC), and the primary visual cortex (V1), as well as in two primate retinal preparations.

We first compared how effectively RMO and other structured stimuli engaged mouse RGC populations relative to WN, and then performed paired comparisons of RF estimation using RMO/RCASE and WN/STA. We then asked which stimulus and analysis factors accounted for differences in RF recovery: we varied the WN square size and frame rate to test the role of spatial and temporal scales, compared moving with static random objects to isolate the contribution of motion, analyzed WN responses with spike-triggered covariance to test for second-order response components not recovered by STA, and compared RMO with cloud stimuli to assess the role of spatial correlations in the display pixel space. Finally, we tested whether RCASE could recover tuning to multiple prespecified object parameters.

Since RMO contains moving contrast edges, we hypothesized that the stimulus may be especially effective at eliciting responses of direction-selective neurons. The NOT receives direct feed-forward input from direction-selective RGCs (DS-RGCs) (Dhande et al., 2013) and, therefore, provides a direct test case of the hypothesis. We evaluated RF recovery separately for direction-selective and non-direction-selective cells to determine whether the advantage of RMO over WN within limited recording time was restricted to direction-selective populations.

## Results

### White noise evokes sparser, weaker, and less reliable spiking than structured stimuli in mouse RGCs

To determine which visual stimuli effectively drive RGC activity, we recorded spiking responses to a battery of light stimuli using high-density microelectrode arrays (HD-MEAs) (Müller et al., 2015); Fig. 1a). The stimulus set included commonly used stimuli for characterizing RGC response properties: a dense binary white noise (WN) stimulus consisting of 100 μm squares, each of which changed its contrast value independently of the others; a full-field chirp stimulus (Baden et al., 2016) (CS); a colorful contrast step stimulus (CCS); a variant of a sparse white noise stimulus that we termed “random static objects” (RSO); and a moving bar stimulus (MB) (Fig. 1b).

**Fig. 1.**
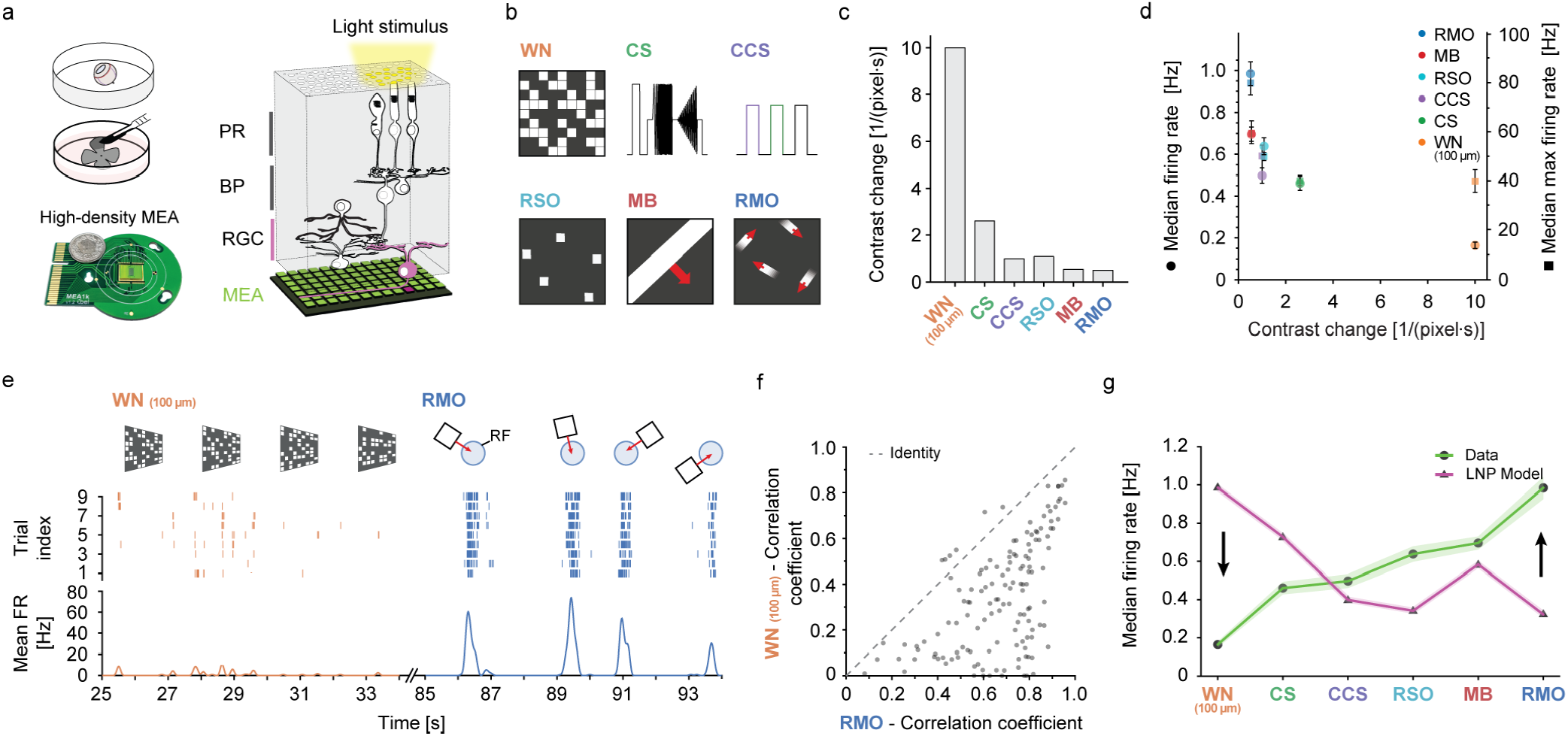
White noise contains frequent contrast changes but evokes sparse, weak, and unreliable retinal responses compared to structured moving stimuli. **a**, Schematic of ex vivo HD-MEA recordings of mouse RGCs with visual stimulation projected onto photoreceptors; simplified retinal circuitry above the array (PR, photoreceptors; BP, bipolar cells; MEA, micro electrode array). **b**, Schematics of the light stimuli used. WN, white noise; CS, full-field chirp stimulus; CCS, colorful contrast step stimulus; RSO, random static objects; MB, moving bar; RMO, random moving objects. **c**, Mean rate of contrast changes per pixel and second for each stimulus. **d**, Contrast-change rate versus population firing statistics (n = 962 RGCs, 3 retinae). Left axis: median time-averaged firing rate. Right axis: median of each cell’s maximum firing rate (computed over the same stimulus epoch; axis scaled for visualization). WN combines the highest contrast-change rate with the lowest median firing rate, while exhibiting a comparatively larger separation between median and maximum firing rate, consistent with rare high-response epochs. Error bars: standard deviation of the median obtained by bootstrap resampling. **e**, Example RGC responses to 9 repetitions (“trials”) of 1 min WN followed by 1 min RMO (9 s excerpts shown; full response in Extended Data Fig. 1). Top: schematic stimulus snapshots; middle: spike raster; bottom: trial-averaged firing rate. **f**, Trial-to-trial reliability: Pearson correlation of firing-rate time series across trials for 146 RGCs (one retina), comparing WN (y-axis) and RMO (x-axis). **g**, Median firing rates across stimuli for recorded RGCs (n = 962, 3 retinae) and LNP model cells (n = 330). Stimuli are ordered by increasing median firing rate in the data. In contrast to the data, LNP responses track stimulus contrast-change rate across stimuli (arrows highlight the discrepancy). Error bars: standard deviation of the median obtained by bootstrap resampling.

We hypothesized that a stimulus containing multiple localized moving objects would strongly engage mouse RGCs. Many mouse RGC types receive strong lateral inhibition and consequently respond poorly to spatially uniform stimuli larger than 1 mm^2^ (Extended Data Table 1), while responding strongly to transient, localized stimuli that match their RF size (Farrow et al., 2013; Roska & Werblin, 2003). Moreover, DS-RGCs, which constitute 20-35% of the entire RGC population (Baden et al., 2016; Chen et al., 2014) in the adult mouse retina, preferentially respond to moving stimuli in specific directions (Dhande et al., 2013, 2019).

Based on these observations, we designed a stimulus consisting of multiple objects moving in random directions, which we termed “random moving objects” (RMO), and added it to the stimulus set (Fig. 1b, bottom right). Each object appeared at a random location on the screen, moved along a straight trajectory for three seconds, and then disappeared. Objects were parameterized by their initial location of appearance and direction of motion.

We then compared how the different stimuli influenced the firing rates of RGCs. As local contrast changes provide the input variation that drives RGC responses (Idrees & Münch, 2020), firing rates evoked by different stimuli cannot be interpreted in isolation: a stimulus may evoke few spikes simply because it provides little local input variation. As a stimulus-level reference, we therefore calculated the average rate of contrast change per pixel for each stimulus. This measure describes how frequently the contrast changes locally, but not how effectively RGCs convert those changes into spikes, which depends on their spatiotemporal filtering and on any selectivity for stimulus features beyond local contrast (Gollisch & Meister, 2010).

The stimuli varied widely in their average rate of contrast change per pixel (Fig. 1c). The WN stimulus featured by far the highest contrast-change rate, as each WN square changed its contrast with a probability of 50% every 50 ms. In contrast, a pixel’s contrast in the MB stimulus changed only twice during a single trial, specifically when the bar edges entered or exited the pixel.

The resulting median RGC firing rates deviated strongly from, and were in fact anticorrelated with the contrast-change rate of the stimuli (Fig. 1d). Although WN exhibited the highest contrast-change rate of all tested stimuli, it elicited the lowest median firing rate. Generally, RGCs showed stronger responses to spatiotemporally sparse stimuli (RSO, MB, and RMO), in which most pixels remained black for extended periods, than to full-field stimuli (CS, CCS) and WN. The two stimuli that induced the highest median firing rates (RMO and MB) were also the only two containing moving contrast edges, while producing comparatively few contrast changes per second. In addition to the median firing rate, we quantified the median of each cell’s maximum firing rate during the same stimulus epoch (Fig. 1d, right axis). For WN, the separation between these two statistics was comparatively large, indicating that responses occurred in rare high-activity epochs despite low average firing.

We next investigated whether the weak firing rates measured in response to the WN stimulus reflected sparse but reproducible responses or unreliable responses throughout the stimulus presentation. To this end, we presented nine repetitions of an identical 2-minute stimulus consisting of one minute of WN followed by one minute of RMO. Many RGCs responded weakly and unreliably to WN but strongly and reliably to the RMO of the same sequence, which we quantified as the trial-to-trial correlation of the firing rate (Fig. 1e,f; Extended Data Fig. 1). The result was robust against changes in the WN square size (Extended Data Fig. 2), or the number of RMO objects simultaneously present per frame (Extended Data Fig. 3a-d) and did not depend on whether the analysis was performed across the entire stimulus duration or 1-s windows (Extended Data Fig. 1f,g; Extended Data Fig. 2c,d). Furthermore, object motion contributed to the strong responses to RMO: the RSO stimulus, which shared the object properties of RMO but lacked motion, evoked systematically lower firing rates than RMO, when the two stimuli were matched either for the average number of objects per frame (Extended Data Fig. 3f) or for the contrast-change rate (Extended Data Fig. 3g).

To better understand why stimuli with a large contrast-change rate can nevertheless evoke weak RGC firing, we used a linear–nonlinear–Poisson (LNP) model (Chichilnisky, 2001; Park & Pillow, 2011; Schwartz et al., 2006) (Extended Data Fig. 4a) adapted from ref. (Jouty et al., 2018). The model linearly filters the stimulus contrast, applies a static nonlinearity, and generates spikes with a Poisson process. We confirmed that the model reproduced the temporal response profiles of common mouse RGC classes to the chirp stimulus on which it was originally parameterized (Jouty et al., 2018), including ON and OFF polarity as well as transient and sustained time courses (Extended Data Fig. 4g). Across stimulus classes, however, the model’s predictions diverged from the data: model firing rates were anticorrelated with the recorded firing rates, and, instead, tracked the stimuli’s contrast-change rates (Fig. 1g; Extended Data Fig. 4h). This mismatch was most pronounced for WN, for which the model predicted high firing rates while recorded RGCs showed the weakest responses, and it was consistent across model cell types (Extended Data Fig. 4i). To verify that this finding did not depend on the parameterization adopted from ref. (Jouty et al., 2018), we fitted LNP models directly to the WN responses of 90 transient ON RGCs from our recordings; the resulting rate estimates were in close agreement with those of the adapted model (Extended Data Fig. 4j). Thus, the weak responses to WN reflected retinal processing not captured by linear contrast encoding, rather than a lack of stimulus input.

### Reverse correlation against stimulus elements (RCASE) for efficient receptive-field estimation

The preceding results highlight a practical system-identification problem. First, a simple LNP model applied to pixel contrast did not account for the large differences in firing rates evoked by different stimulus classes. Second, stimuli designed for classical linear system identification did not necessarily elicit strong and reliable responses within experimentally practicable recording times. Therefore, we asked whether the strong, reproducible responses evoked by locally structured stimuli could be leveraged to estimate RFs and tuning properties more efficiently.

We first outline linear RF estimation using STA on the responses to the WN stimulus and then show how the same reverse-correlation logic can be applied to a stimulus-element representation to estimate spatial and temporal response kernels as well as parameter-tuning functions (Fig. 2). A key advantage of WN is its spatial homogeneity: since all pixels follow the same statistics, each neuron experiences comparable stimulus statistics regardless of where its RF lies, which enables parallel characterization of large populations.

**Fig. 2.**
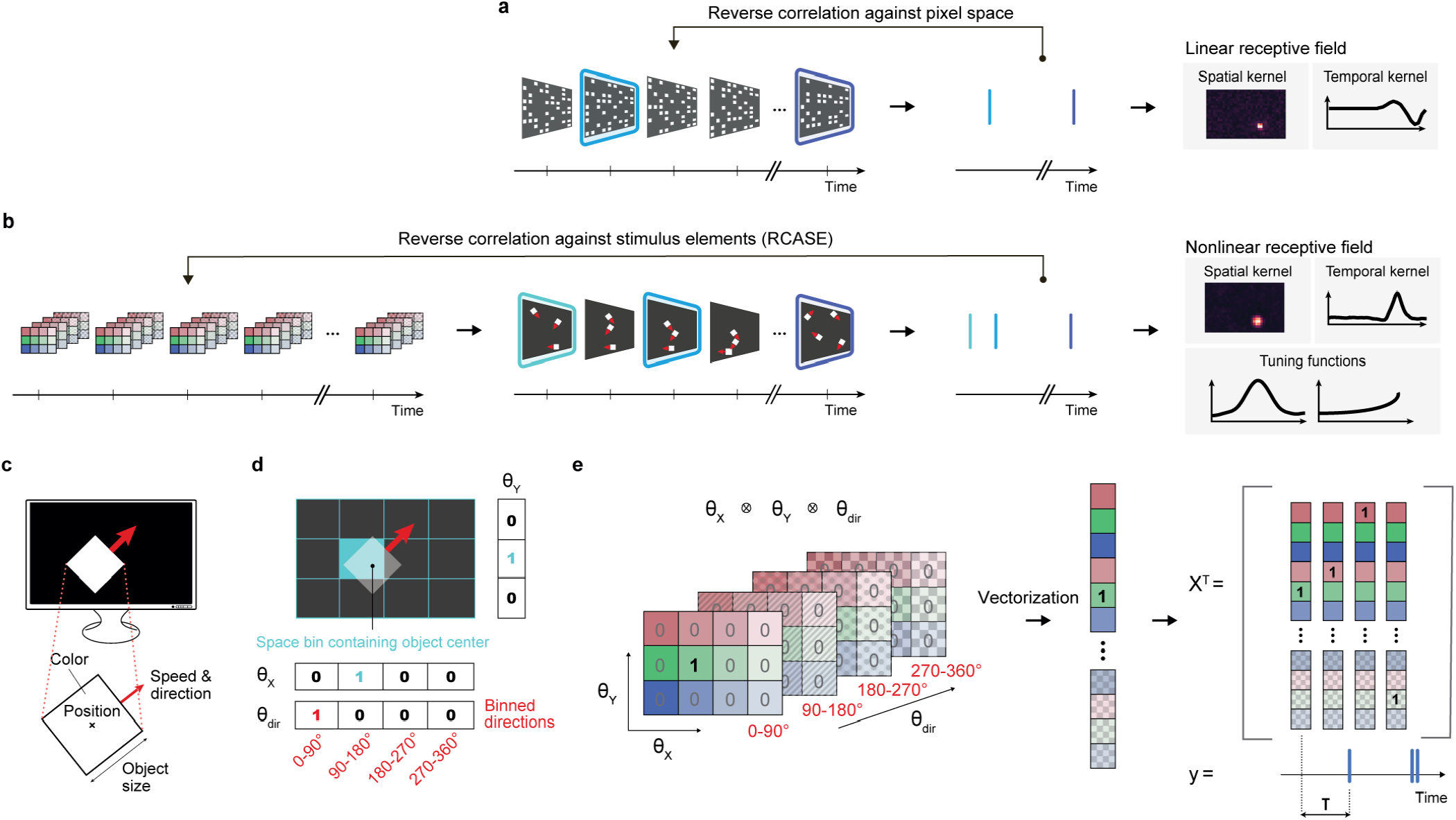
RCASE generalizes spike-triggered averaging by reverse correlation of spikes against structured stimulus elements. **a**, Method illustration: estimation of linear RFs (spike-triggered averages) by reverse correlation of the neural response against a white-noise stimulus. Spikes select stimulus frames preceding each spike; averaging the selected stimulus segments yields a linear spatiotemporal RF (spatial and temporal kernels). **b**, RCASE for RMO. Left: a sparse multidimensional stimulus-element tensor encodes the parameters of the stimulus elements (e.g., position on the display) for each movie frame. A series of stimulus elements encodes the trajectory of a moving object, and multiple objects (multiple stimulus elements) can be visible at the same time. Middle: corresponding stimulus movie and neural response (blue marks denote spike times; red arrows illustrate motion direction and are not part of the stimulus). Right: schematic of the output obtained by reverse correlation of the response to the stimulus-element representation, yielding spatial/temporal kernels and parameter tuning functions. **c**, Illustration of stimulus elements on the display and example parameters of a stimulus element. **d**, Simplified encoding example. A stimulus element is parameterized by x- and y-location and direction of movement. After binning each parameter dimension, the presence of an element is encoded by a binary vector with a single active entry for each parameter (one-hot encoding). **e**, Construction of the stimulus matrix used for reverse correlation. The outer product of the one-hot vectors defines a high-dimensional but sparse tensor encoding a single stimulus element in one frame (left). To encode multiple stimulus elements in a single frame, the corresponding tensors are summed. The resulting tensor is vectorized (middle) to yield a row of the stimulus matrix X (right), which is reverse correlated against the neural response y.

In WN analysis, linear RFs can be estimated by reverse correlation of the neural response with the stimulus contrast (Chichilnisky, 2001; D. Ringach & Shapley, 2004; Schwartz et al., 2006). Reverse correlation can be formulated as a linear regression problem (Park & Pillow, 2011), where the regressor is a stimulus matrix, the rows of which contain the WN square contrasts in each time bin, and where the response variable is the spike count per time bin. The ordinary least-squares estimator yields the maximum-likelihood estimate of the RF, or equivalently, the whitened spike-triggered average (Park & Pillow, 2011). Repeating the estimation for different response lags gives a spatiotemporal RF that can be decomposed into spatial and temporal kernels (D. Ringach & Shapley, 2004) (Fig. 2a). Under idealized assumptions (*e*.*g*., Gaussian WN), the stimulus distribution is spherically symmetric in stimulus space, which supports unbiased linear-filter estimates (Chichilnisky, 2001); in practice, binary WN is often used to increase firing rates, which may introduce deviations from these ideal assumptions and lead to biases (Paninski, 2003).

We designed the RMO stimulus to retain a similar form of spatial homogeneity by drawing object parameters, including position and motion direction, from uniform distributions across the display. Unlike WN, however, the resulting pixel intensities were strongly structured and correlated in space and time. Consequently, reverse correlation with pixel contrast yields biased linear-filter estimates, and correcting for the stimulus correlations is numerically ill-posed because of collinearities in the pixel-space stimulus matrix (i.e., a singular or ill-conditioned Gram matrix).

To address this issue, we developed “reverse correlation against stimulus elements” (RCASE) (Fig. 2b). Its central idea is to perform reverse correlation in a stimulus-element representation rather than in pixel space. The estimated kernels are, therefore, linear functions of the stimulus-element variables rather than of the pixel contrast, and generally nonlinear with respect to display pixels. RCASE is designed to efficiently characterize hypothesized low-dimensional features (*e*.*g*., parameterized moving objects), rather than to perform unconstrained feature discovery in arbitrary high-dimensional stimulus spaces.

For the RMO stimulus, a stimulus element describes the presence of an object with a specific set of parameters within a single stimulus frame (Fig. 2c). For clarity, we considered a simplified parameterization consisting of x- and y-position *θ*_*x*_, *θ*_*y*_ and motion direction (*θ*_*dir*_). Each parameter axis was discretized into bins, and an element was assigned to one bin per parameter (Fig. 2d). The element was then encoded by one-hot vectors for each parameter, whose outer product forms a sparse stimulus-element tensor with a single active entry at the element’s coordinates in the parameter space (Fig. 2d,e). If multiple elements were present in a frame, their tensors were summed.

The resulting tensor was vectorized, and repeating this procedure across frames yielded the RCASE design matrix *X* (Fig. 2e). This matrix was then reverse-correlated against the neural response *y* . Because object parameters were drawn independently from uniform distributions, the covariance of *X* within a frame was approximately diagonal, so that this stimulus representation was effectively free of the instantaneous spatial correlations present in the pixel-space representation. This spatial independence is analogous to that of WN squares. The design matrix thus supported well-conditioned least-squares estimation at each temporal lag. In contrast to WN, however, the RMO stimulus contains temporal correlations across consecutive frames, because each object persists along a deterministic trajectory (see “Temporal correlations in the RMO design matrix” in the Discussion). These temporal correlations must be considered when interpreting kernel shapes across lags.

Beyond these statistical properties, WN/STA and RMO/RCASE describe the neural response with respect to different stimulus spaces. For neurons, the response of which contains a non-vanishing linear component in pixel space, WN/STA will converge given sufficient data. Under limited experimental time, however, the choice of stimulus can strongly affect estimation efficiency. We, therefore, hypothesized that a stimulus evoking strong and reliable responses, analyzed in the more abstract stimulus-element space, would allow RFs to be recovered within a practicable recording time (here, 20 min) while remaining directly interpretable as tuning to the prespecified element parameters.

### Strong, informative responses to RMO enable rapid receptive-field estimation within a few minutes

For each RGC, we used responses to the RMO stimulus to estimate an RF with RCASE (RMO-RF) and responses to the WN stimulus to estimate an RF with STA (WN-RF) (n=962 RGCs, 3 retinae). We identified RGCs with co-localized RMO-RFs and WN-RFs, as well as RGCs exhibiting exclusively RMO-RFs (Fig. 3a). To quantify the quality of the spatial RF obtained with each stimulus-analysis combination, we defined an RF quality index (Baden et al., 2016) *RF*_*QI*_ as the fraction of variance explained by a two-dimensional Gaussian fit to the cell’s spatial RF. An *RF*_*QI*_ of 1 indicates that the fit completely explains the spatial RF, while an *RF*_*QI*_ close to 0 indicates a noisy spatial RF. We defined cells with an *RF*_*QI*_ > 0.3 to have a discernible spatial RF. We found that while only 44.3% of RGCs displayed discernible WN-RFs after 20 min of stimulation, we obtained discernible RMO-RFs for 97.9% after the same duration (Fig. 3b,c).

**Fig. 3.**
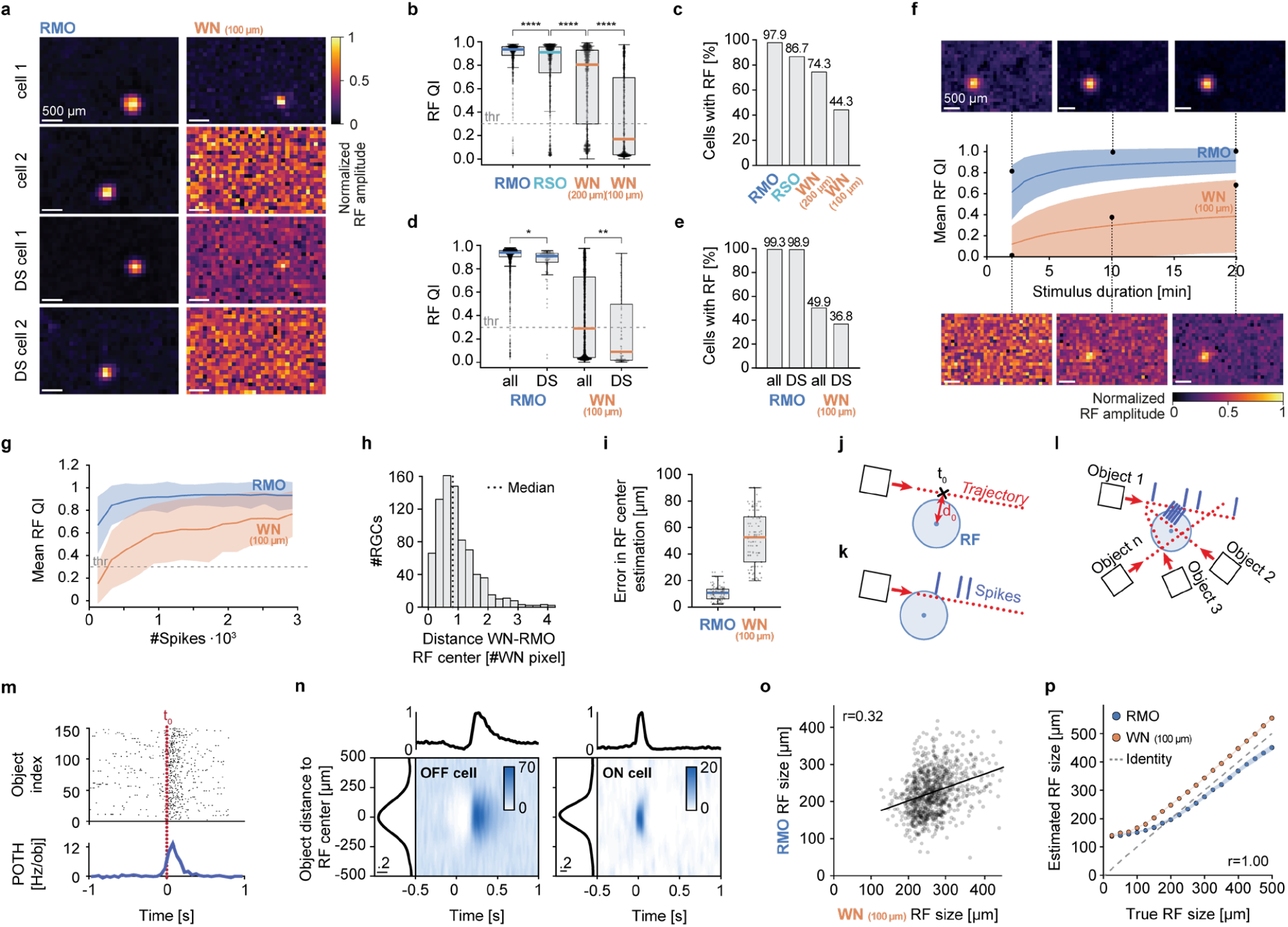
Comparison of RF properties estimated with RMO and WN stimuli in the mouse retina. **a**, Single cell examples of spatial RFs for Non-DS (rows 1&2) and DS-Cells (rows 3&4). Color indicates RF strength from low (black) to high (white). **b**, RF quality index *RF*_*QI*_ for the recorded cells (n=962, 3 retinae) for the RMO, RSO, and WN stimulus (20 min each) with 100 μm (WN) and 200 μm pixel size (WN 200 μm); (**** P < 0.0001, Wilcoxon signed-rank test). **c**, Percentage of all recorded RGCs with an *RF*_*QI*_ > 0.3 for the different stimuli. **d**, *RF*_*QI*_ for recorded RGCs (n=1871, 4 retinae) and for DS-RGCs (n=96) for the RMO and WN stimulus; (*P < 0.05, **P < 0.01, Mann–Whitney U test). **e**, Percentage of all recorded RGCs and DS-RGCs with an *RF*_*QI*_ > 0.3 for WN and RMO stimuli. **f**, Mean *RF*_*QI*_ as a function of stimulus duration; shaded area: standard deviation. Examples of estimated spatial RFs are shown as images for each stimulus at a stimulus duration of 2, 10, and 20 minutes. **g**, Mean *RF*_*QI*_ as a function of spike count; shaded area: standard deviation. **h**, Distance between RMO- and WN-RF centers of RGCs with *RF*_*QI*_ > 0.3 for both RMO and WN (n=793, 3 retinae). **i**, Difference between estimated and true RF center location for LNP-model RGCs (n=110). **j-l**, Illustration of the peri-object time histogram (POTH) method for the RMO stimulus. **j**, For each moving object and each neuron, the closest-approach time *t*_0_ is determined as the time point when the object was closest to the neuron’s RF center with closest approach distance *d*_0_. **k**, The object is assigned a shifted version of the neuron’s spike train, aligned to *t*_0_. **l**, For each neuron, the process is applied to all objects that passed the neuron’s RF. **m**, Top: Raster plot of an example neuron’s responses to multiple objects (rows) aligned to *tt*_0_. Bottom: Average firing rate over all objects (POTH). **n**, Estimation of RF size utilizing the POTH method. Distance-time histograms for an OFF and an ON cell. Each row is a POTH for all objects within a specific distance bin (y-axis). Colors indicate Hz per object. Histograms above and on the side: Time and distance components, obtained by singular value decomposition. **o**, Comparison of RMO and WN RF sizes for RGCs with *RF*_*QI*_ > 0.3 for both RMO and WN (n=913, 4 retinae). Black line: linear regression fit (r=0.32). **p**, Comparison of model (true) and estimated RMO- and WN-RF size for a population of LNP model cells (n=500). Each point is the average over 25 LNP model cells with the same model RF size; shaded area: standard deviation. Dashed line: identity function.

As the objects in the RMO stimulus (200 *µm* edge length) were larger than the WN squares (100 *µm* edge length), we investigated whether this size mismatch could account for the observed difference in RF estimation. Increasing the WN square edge length to 200 *µm* increased the fraction of RGCs with a discernible WN-RF to 74.3%. However, a substantial number of RGCs still showed no WN-RFs (Fig. 3b,c). Additionally, this increase in size came at the cost of reduced spatial resolution of the WN-RFs.

An alternative to enlarging the WN squares is to introduce spatial correlations into the noise itself. We, therefore, also presented a spatially correlated “cloud” stimulus (Shi et al., 2019), with stimulus power concentrated at low spatial frequencies, together with a contrast-normalized version of the same stimulus (n = 287 RGCs, 2 retinae; Extended Data Fig. 5g–i). After 20 min of stimulation, the cloud stimulus yielded discernible RFs for 19.9% of RGCs, no more than binary WN in the same recordings (20.6%), whereas the normalized-cloud stimulus increased this fraction to 62.4%. RMO/RCASE yielded discernible RFs for 99.3% of the same cells (Extended Data Fig. 5h,i). Thus, spatial correlations in pixel space improved WN-based RF estimation only when contrast was restored, and even then fell well short of RMO/RCASE.

To isolate the contribution of object motion to RF estimation, we compared the RMO stimulus with the RSO stimulus (Fig. 1b), which contained static white objects of the same size. In the RSO stimulus, multiple objects with an edge length of 200 *µm* appeared simultaneously at random locations and remained visible for three stimulus frames (50 ms) before being replaced by a new set of objects at different random locations. The RSO stimulus thus resembled a sparse white noise stimulus with the same refresh rate (20 Hz) as the WN stimulus. As with the RMO stimulus, we encoded the RSO stimulus as a matrix representing the presence or absence of stimulus elements and analyzed the neural responses with RCASE. We found that 86.7% of RGCs had a discernible RSO-RF (Fig. 3c), substantially more than with 100 *µm* WN (44.3%) but fewer than with the RMO stimulus (97.9%). Thus, for a subset of RGCs, an RF could be identified exclusively with the RMO stimulus, indicating that object motion was necessary for RF estimation in these cells within 20 min of stimulation.

Because the RMO stimulus contains moving contrast edges, we asked whether its advantage over the WN stimulus was especially pronounced for DS-RGCs. We, therefore, identified DS-RGCs using the MB stimulus and compared RF estimation in the DS subset with that in the full RGC population. For this analysis, we compared 1,871 RGCs (4 retinae) with a subset of 96 DS-RGCs (single-cell examples in Fig. 3a, bottom rows). We found notable differences for WN-RFs between the two populations, both in the median *RF*_*QI*_ (0.29 for the full population vs. 0.10 for DS-RGCs; Fig. 3d) and in the fraction of RGCs with a discernible WN-RF (49.9% vs. 36.8%; Fig. 3e). For RMO-RFs, however, the two populations were nearly indistinguishable, both in the median *RF*_*QI*_ (0.94 vs. 0.93; Fig. 3d) and in the fraction of RGCs with a discernible RMO-RF (99.3% vs. 98.9%; Fig. 3e).

The quality of a WN-RF estimate depends on the number of spikes recorded during stimulation (Schwartz et al., 2006). Given that RGCs responded more strongly to the RMO than to the WN stimulus, we investigated how rapidly we could estimate RMO-RFs compared to WN-RFs. We estimated WN-RFs and RMO-RFs as a function of stimulus duration in a range of 2 to 20 min in 1-min steps. While it took 14 min of stimulation to identify 90% of the identifiable WN-RFs, the same percentage of RMO-RFs could be detected within just 3 min (Fig. 3f). Considering the limited experimental time, the ability to rapidly estimate RMO-RFs, therefore, presents a substantial advantage.

To investigate whether the greater efficiency of RMO/RCASE was explained solely by the higher firing rates in response to the RMO stimulus, we evaluated RF quality as a function of spike count rather than stimulus duration. Notably, RMO/RCASE produced high-quality RF estimates with substantially fewer spikes than WN/STA, with RF quality quickly reaching a plateau (Fig. 3g). Thus, the greater efficiency of RMO/RCASE could not be explained solely by the higher firing rates evoked by the RMO stimulus.

Taken together, these results show that RMO/RCASE yielded discernible spatial RFs for substantially more mouse RGCs and required less recording time than WN/STA. Notably, the time advantage did not stem from higher firing rates alone: RF quality saturated at far lower spike counts, suggesting that individual spikes, evoked by the RMO stimulus, carried more information about a cell’s RF than those evoked by the WN stimulus. The advantage of RMO/RCASE was particularly apparent in DS-RGCs, for which WN/STA performed poorly, but was not restricted to that population.

### Comparison of spatial receptive-field estimates obtained with RMO/RCASE and WN/STA

RMO/RCASE and WN/STA characterize responses in different stimulus representations, and their estimates of spatial extent can therefore differ. We first compared the estimated RF centers using both approaches and then examined how object size, object motion, and temporal correlations affect the apparent spatial extent of RMO-RFs.

To compare RMO-RFs and WN-RFs, we examined the accuracy of the RF center estimation of 793 RGCs (3 retinae) with both discernible WN- and RMO-RFs. After upsampling and smoothing, we defined each RF center as the spatial location of the peak inside the RF. WN-RF and RMO-RF centers co-localized with a median distance of 82 μm, smaller than the 100-μm spatial resolution of the WN stimulus (Fig. 3h) which was limited by the size of the WN squares. In contrast, RMO objects moved between pixels of the display, which meant that the spatial resolution of the RMO stimulus was constrained only by the resolution of the display. To directly assess localization accuracy, we conducted an analysis utilizing LNP models, for which the correct spatial position of their RF was known. It demonstrated markedly lower errors in estimating the RF centers with RMO-RFs compared to WN-RFs (Fig. 3i). The rapid and accurate localization of RMO-RFs may thus provide a substantial methodological advantage when a precise centering of the stimuli on RFs is required.

The spatial extent of the WN-RF is an important parameter for identifying RGC subtypes and mosaics (Baden et al., 2016; Chichilnisky & Kalmar, 2002). Since the objects of the RMO stimulus have spatial extents themselves, RGC responses may occur well before these objects are centered on the RGC RFs and potentially persist when the object leaves the RF center. Furthermore, unlike spatiotemporally uncorrelated WN, RMO objects were present over multiple frames, which induces temporal correlations in the RCASE stimulus design matrix and must be considered when interpreting the resulting RFs. These issues raise the question of how to interpret the apparent size of the RMO-RF, as it depends on both the size and speed of the moving objects.

To estimate RF sizes in the presence of temporal correlations, we developed a complementary method for analyzing the RGC responses to the RMO stimulus, which we refer to as peri-object time histogram (POTH), in reference to the peri-stimulus time histogram (Gerstein & Kiang, 1960). We first identified a cell’s RF center by locating the maximum of the spatial RF using RCASE. For each object, we tracked the distance between its center and the RGC’s RF center throughout its 3-second lifetime. We then determined each object’s minimum distance to the RF center, d_0_, and time of closest approach, t_0_ (Fig. 3j). Each object was then assigned a shifted version of the neuron’s spike train, aligned to t_0_ (Fig. 3k,l). We used aligned spike trains to construct a POTH (Fig. 3m), grouped objects according to their minimum distance from the RF center in 50-μm bins, and then computed a POTH for each group, yielding a two-dimensional response distance-time histogram for each RGC (Fig. 3n). Using this approach, we estimated the distance between an object’s center and the RGC’s RF center at which RGCs ceased to respond. This served as an effective proxy for the RGC’s RF size. POTH-based RF-size estimates positively correlated to WN-RF sizes (*r* = 0.32, *p* < 10^−22^), although with considerable variability (Fig. 3o).

To understand whether this variability reflected retinal processing, where object motion may influence the spatial extent of an RF (Deny et al., 2017), we used an LNP model. We systematically varied the RF sizes of the LNP model, computed model responses to the WN and RMO stimuli, and estimated the resulting RMO- and WN-RF sizes. Both methods produced biased RF-size estimates when the underlying RFs were smaller than the WN square size or RMO object size. For larger RFs, however, the results of both methods showed a perfect correlation between the estimated RMO- and WN-RF sizes (r=1.0; Fig. 3p). This finding suggested that if the assumptions underlying the WN analysis were fulfilled by the neuronal system, *i*.*e*., if the cells responded linearly to the stimulus, both methods would agree on the RF sizes. Therefore, the discrepancy between the RF-size estimates obtained with the two methods is unlikely to arise from the analysis itself but more likely reflects properties of the underlying neural responses.

### Estimation of multiple tuning functions in parallel from RMO-RFs and their temporal dynamics

ON- and OFF-RGCs respond preferentially to contrast changes from dark to light and from light to dark, respectively. Consequently, their WN-RFs display distinct temporal dynamics revealing their ON or OFF preference (Chichilnisky & Kalmar, 2002). In contrast, ON-OFF RGCs respond to both positive and negative contrast changes, a response property that cannot generally be captured by a single linear function of the pixel contrast. For balanced ON-OFF responses, the contributions of opposite contrast polarities cancel in the linear STA, preventing recovery of the underlying RF (Schwartz et al., 2006). When responses to the two polarities are unbalanced, the WN-RF instead tends to reflect the temporal dynamics of the dominant response, obscuring the underlying ON-OFF response structure. Covariance-based approaches, such as spike-triggered covariance (STC), can recover multi-feature or symmetric nonlinear RF structures from neural responses to WN (Cantrell et al., 2010; Pillow & Simoncelli, 2006; Rust et al., 2005; Schwartz et al., 2006). However, the number of spikes required for STC analysis scales quadratically with stimulus dimensionality (Ahn et al., 2020), making full-resolution WN impracticable to use unless the RF location is already known and dimensionality reduction can be applied. Consistent with this limitation, we show that STC recovered the expected RF structure in modeled ON-OFF cells but performed poorly on our experimental WN recordings at full resolution (Extended Data Fig. 5). Even when provided with RCASE-estimated RF locations for dimensionality reduction, STC recovered significant RF structure in only ∼18% of the cells with 100-µm WN and ∼44% with 200-µm WN (Extended Data Fig. 5d).

Next, we investigated whether the temporal dynamics of RMO-RFs could distinguish between ON, OFF, and ON-OFF RGCs. RMO-RF temporal kernels differed across the three response classes; ON cells exhibited earlier response peaks than OFF cells, whereas single versus multiple peaks distinguished ON and OFF cells from ON-OFF cells (Extended Data Fig. 6a). The timing of the RMO response peak was strongly negatively correlated with an ON-OFF index that was independently determined from contrast-step responses in the same cells (r=-0.73; Extended Data Fig. 6b). Thus, RMO-RF temporal dynamics captured response properties that distinguish ON-, OFF- and ON-OFF RGCs.

The single WN/STA kernel estimated here does not directly provide a direction-tuning curve. In RCASE, in contrast, stimulus elements can be indexed by motion direction, allowing direction tuning to be estimated directly from the RMO responses. Directional tuning functions can be derived using RCASE as follows. We estimated RMO-RFs by binning the angular direction of movement of each object, *θ*_*dir*_, into equally sized bins and calculated the temporal kernel for each direction bin separately. Fig. 4e,f illustrates this analysis for an example non-DS-RGC and Fig. 4g,h for an example DS-RGC. The process yielded a two-dimensional direction-time response matrix (Fig. 4i), revealing clear direction selectivity for DS-RGCs (Fig. 4g,h,i). We determined direction-tuning curves of 55 DS-RGCs (7 retinae) that showed direction-selective responses to both moving bar and RMO stimuli. Both stimuli yielded similar tuning curves (Fig. 4j). We then determined the preferred direction for each stimulus by a von object-pairs Mises fit to the respective direction tuning curves and found that they were strongly correlated (circular correlation coefficient (Jammalamadaka & Sengupta, 2001): 0.8; Fig. 4k). This finding showed that tuning functions to the stimulus element parameters were robustly estimated for the recorded DS-RGCs.

**Fig. 4.**
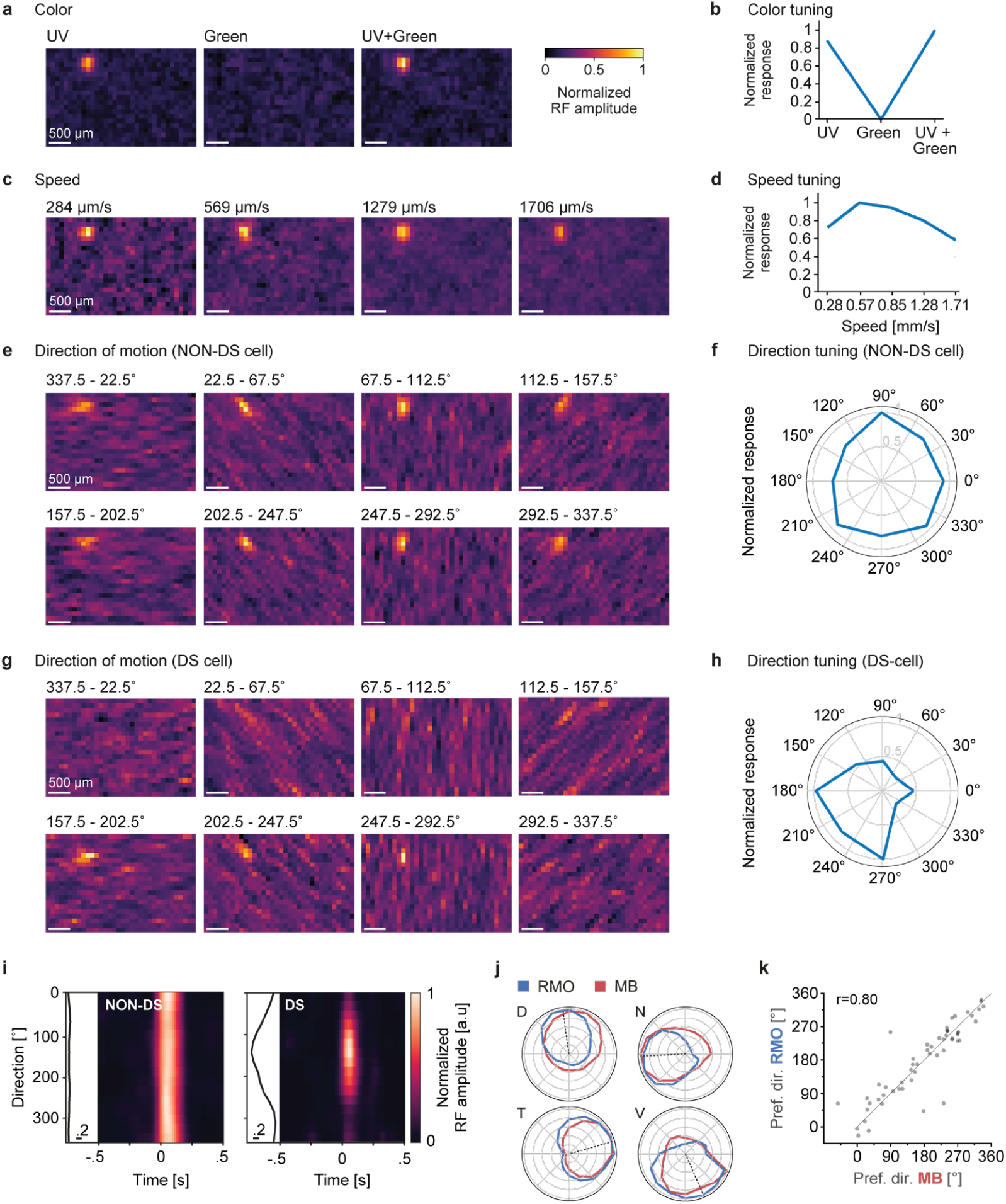
Parallel estimation of spatial receptive fields and object-parameter tuning. **a**, Spatial RFs of an example non-direction-selective (non-DS) retinal ganglion cell, estimated separately for the three object colors of RMO stimulus (UV, green, and UV + green). Color indicates the normalized RF amplitude on a common scale across the three colors. Scale bar: 500 μm. **b**, Max-normalized color tuning curve of the same cell. **c**, Spatial RFs of the same cell, estimated separately for five object speeds. Here results for four speeds are shown. Color indicates the normalized RF amplitude on a common scale across the four speeds. Scale bar: 500 μm. **d**, Max-normalized speed tuning curve of the same cell. **e**, Spatial RFs of the same cell, estimated separately for eight object movement directions. Color indicates the normalized RF amplitude on a common scale across the eight directions. Scale bar: 500 μm. **f**, Max-normalized direction tuning curve of the same cell. **g**, Spatial RFs of an example direction-selective (DS) retinal ganglion cell. Color indicates the normalized RF amplitude on a common scale across the eight directions. Scale bar: 500 μm. **h**, Max-normalized direction tuning curve of the same DS cell. **i**, Direction–time matrices for an example non-DS retinal ganglion cell (left) and an example DS retinal ganglion cell (right). Each row is the RMO temporal kernel estimated for objects moving within one of 16 equally sized direction bins (22.5° wide, spanning 360°). Color indicates the temporal-kernel amplitude, normalized to the maximum of each direction–time matrix. The direction tuning curve, obtained by singular value decomposition of the matrix and then normalized by the maximum, is shown at the left of each panel. **j**, Direction tuning curves of an example DS retinal ganglion cell from each of the four retinal quadrants (dorsal, nasal, temporal, and ventral), giving each cell’s max-normalized response across 16 movement-direction bins (22.5° wide) estimated with the RMO (blue) and the moving-bar (MB, red) stimulus. Dashed line: preferred direction determined by a von Mises fit to the RMO direction tuning curve. **k**, Preferred movement direction of direction-selective RGCs estimated from the RMO stimulus versus the moving-bar (MB) stimulus (n=55, 7 mouse retinae). For each cell, the preferred direction was determined separately for each stimulus by a von Mises fit to its direction tuning curve. Each gray dot is one RGC; dashed line: identity. The preferred directions estimated from the two stimuli were strongly correlated (circular correlation coefficient r = 0.80).

In a separate set of experiments (n=795, 3 retinae), we presented a modified RMO stimulus containing objects of variable speed and color to assess if tuning functions for multiple parameters could be estimated in parallel (Fig. 4a-d). We performed hierarchical clustering of RGCs based on their RMO-RF temporal kernels and tuning functions to the object distance, speed, and color, identifying 15 clusters (Extended Data Fig. 6c). The resulting clusters exhibited distinct combinations of temporal response properties and tuning to object distance, speed, and color (Extended Data Fig. 6d,e). Thus, RMO/RCASE enabled parallel estimation of multiple response properties across the recorded population upon applying a single stimulus, and there is no need to present multiple specific stimuli to estimate individual response parameters or to pre-align the stimulus with the RF locations.

### Responses to successive stimulus elements reveal temporal interactions in mouse retina

Using the RMO stimulus, multiple objects can move in short succession over an RF. We investigated whether a neuron’s response to an object was influenced by a second object that passed its RF shortly beforehand. Furthermore, we wanted to estimate if the polarity of the objects, i.e., whether they were bright or dark, determined this influence. For that purpose, we presented a modified RMO stimulus consisting of a grey background and moving black and white objects. For each identified RF, we selected objects that passed the RF center at a maximal distance of 100 μm, thus traversing the RGCs’ RF. We then searched for pairs of objects that moved sequentially over the RF, separated by the time lag Δ*t* (Fig. 5a), and binned object pairs according to their inter-object time lags in bins of 100 ms. Finally, object pairs were grouped into four possible color combinations: white-white, white-black, black-white, or black-black. For each pair, we defined the first and second objects as A and B, respectively, and aligned the spike train to the time at which object B was closest to the RF center: *t*_0,*B*_. We then calculated the POTH over the object pairs, resulting in a 2D-histogram for each color combination (Fig. 5b,c; Extended Data Fig. 7). Averaging POTHs for the white-white condition across 105 ON RGCs (one retina) revealed an increased response for Δ*t* = 0*s*, which was to be expected given that two white objects were simultaneously present within the RF (Fig. 5b). In contrast, for large inter-object lags, such as Δt > 0.8 s, the response to object B was largely unaffected by the preceding object A. Notably, we observed suppression of the response to object B at a small Δ*t* ≈200 ms. We hypothesized that the suppression reflected an inhibitory surround mechanism.

**Fig. 5.**
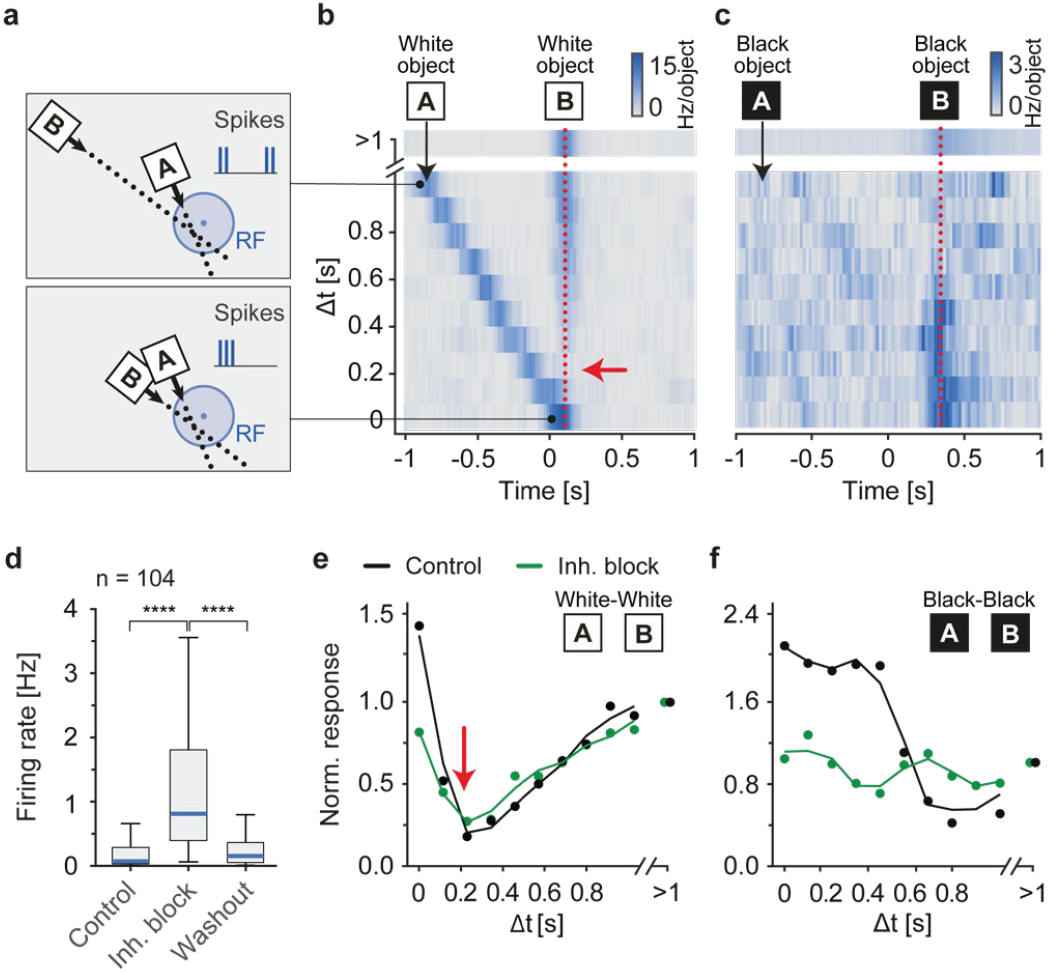
Second-order stimulus interactions reveal nonlinear neural responses. **a**, Schematic of the analysis of the neural response to pairs of objects that pass a neuron’s RF with a specific temporal distance (Δ*t*). Top: Object A moves before object B over the cells’ RF. Bottom: Object A and object B move over the RF at similar times, i.e. they have similar closest approach times *t*_0,*A*_ ≈ *t*_0,*B*_ and Δ*t* = *t*_0,*B*_ − *t*_0,*A*_ ≈ 0. **b**, Population-averaged (n=105 ON-Cells) 2D-Histogram of responses to pairs of white objects with different Δ*t* (responses to other combinations of object colors in Extended Data Fig. 7). Responses are aligned with respect to object B’s closest approach time, i.e. *t*_0,*B*_ = 0. Red dotted line highlights the time bin shown in e (control) in isolation. Red arrow indicates suppression of the response to object B. **c**, Same as b, for black object pairs. Red dotted line highlights the time bin shown in f, (control) in isolation. **d**, Firing rate over 40 min of RMO stimulation of ON-cells (n=105) in three conditions: control, pharmacological block of amacrine-cell mediated inhibition (strychnine, gabazine, picrotoxin), and washout; (****P < 0.0001, Wilcoxon signed-rank test). **e**, Response to object B for two white objects as a function of Δ*t* normalized to the response for Δ*t* > 1 *s*. Red arrow indicates suppression of the response at Δ*t* = 0.25*s*. **f**, Same as e for a pair of black objects.

To test the contribution of inhibitory retinal circuitry, we administered a pharmacological cocktail consisting of strychnine (1 μM), gabazine (5 μM), and picrotoxin (100 μM) to block glycine and GABA_A_-mediated inhibition (Yan et al., 2020). Despite the strong increase in the population mean firing rate after pharmacological perturbation (Fig. 5d), the suppression persisted (Fig. 5e). This finding suggests that the suppression was a result of local retinal computations and not of long-range inhibition. A model proposed previously (Idrees et al., 2022) provides a potential explanation of such a suppressive mechanism without inhibitory interactions, through an interplay of cone dynamics and threshold nonlinearities. In the model, relatively slow cone dynamics cause the responses to successive objects to overlap temporally. This has an effect similar to temporarily increasing the threshold of the threshold nonlinearities and, therefore, block the response to the second object. Our results indicate that such a mechanism most likely also plays a role in the encoding of successive movements of contrast edges across the RFs of RGCs.

Additionally, when two black objects successively traversed the RFs of ON RGCs, the second object in contrast elicited an enhanced response (Fig. 5c,f), indicating that the response to a dark object depended on the recent passage of another dark object across the RF. Contrary to the persistent suppression for successive bright objects, blocking inhibition abolished the enhanced response to the second dark object. Consequently, our findings imply disinhibition as a key mechanism underlying the pronounced spike response in ON RGCs following the passage of dark objects through their RFs. Beyond first-order kernels and tuning functions, the stimulus-element framework thus gives experimental access to interactions between successive stimulus elements and, as shown here, to the circuit mechanisms underlying them.

### Receptive-field estimation across the early visual system

To test whether the advantage of RMO/RCASE over WN/STA generalized to populations with different response properties, we compared RF estimation across four visual structures of the mouse early visual system: retina, superior colliculus (SC), nucleus of the optic tract (NOT), and primary visual cortex (V1). Recordings from SC, NOT, and V1 were performed using Neuropixels probes (Steinmetz et al., 2021) in anesthetized head-fixed mice passively exposed to the light stimuli.

We first compared the strength of the stimulus-evoked responses across visual areas. Neurons across all visual structures responded robustly to the RMO stimulus (Fig. 6a–e; Extended Data Fig. 8a, 9a). However, the relative response strength evoked by different stimuli varied across visual structures. In mouse retinal DS-RGCs, the CS elicited stronger responses than WN, although responses to both stimuli remained substantially weaker than those to RMO (Fig. 6a). SC neurons exhibited more comparable response strengths across stimuli (Fig. 6b). In contrast, NOT neurons responded weakly to CS despite exhibiting robust responses to RMO (Fig. 6c). Furthermore, the differences in firing rate evoked by RMO and WN were less pronounced in NOT and SC, compared to retinal DS-RGCs. Finally, V1 neurons responded strongly to all stimulus classes, including the CS (Fig. 6d).

**Fig. 6.**
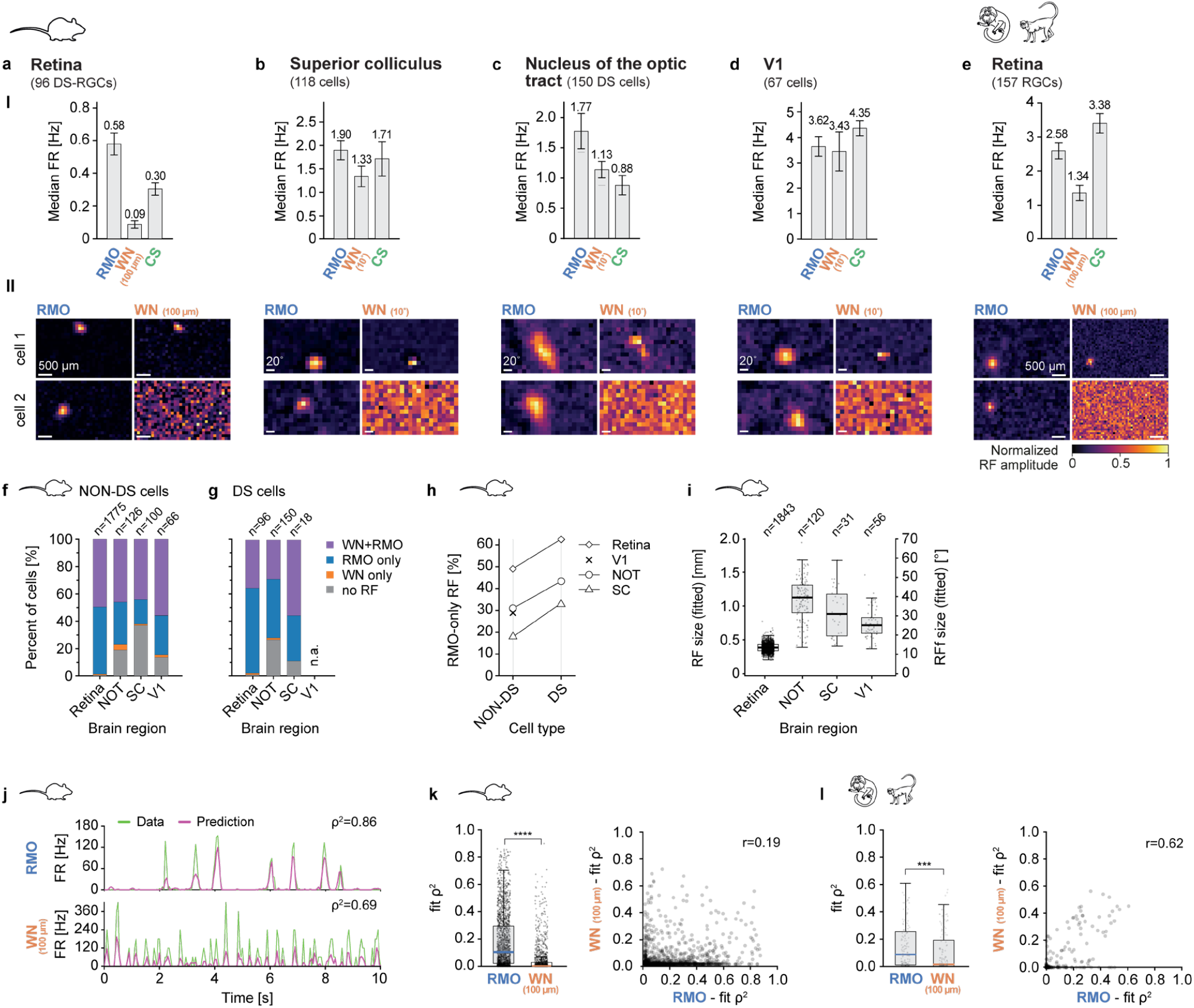
Performance of RMO/RCASE and WN/STA in different mouse visual areas and primate retina. **a-e**, Median firing rate across cell populations for different light stimuli and brain areas; **a**, Mouse DS-RGCs (n=96, 4 retinae); **b**, Mouse superior colliculus (SC) (n=118, 3 mice); **c**, DS-Cells in the mouse nucleus of the optic tract (NOT) (n=150, 7 mice); **d**, Non-DS cells in mouse V1 (n=67, 4 mice); The DS population was too small to be reported. **e**, Primate RGCs (n=157, 1 marmoset retina, 1 macaque retina – analysis per species in Extended Data Fig. 9). Error bars: standard deviation of median firing rates, obtained through bootstrap resampling. II, Spatial RFs for RMO and WN for two example cells from same cell populations in I. **f-g**, Percentage of RFs reconstructed with different methods across mouse cell populations for NON-DS and DS cells. Note: we do not report DS cells in V1 as the recorded population was too small. **h**, Percentage of RMO-reconstructed RFs across mouse NON-DS and DS cell populations. **i**, RF sizes across mouse cell populations. **j**, Response prediction of an LN-model fitted to example mouse RGCs. Top: RGC response and LN-model prediction for an example mouse RGC for the RMO stimulus. Shown are 10s of the response. Bottom: Same as above for the WN stimulus. **k**, Comparison of WN and RMO prediction performance (10-fold cross-validated squared correlation coefficient on the test set, *ρ*^2^) for the mouse retina (n=1346, 4 retinae). Left: Box plot; Right: Scatter plot (****P < 0.0001, Wilcoxon signed-rank test). **l**, Same as k for primate retina; (***P < 0.001, Wilcoxon signed-rank test).

Using the same WN square size in NOT as in the retinal recordings yielded few discernible WN-RFs (12.5%; Extended Data Fig. 10b,c). We, therefore, increased the WN square size to a visual size corresponding to 288 µm on the retina (or 10° in visual degrees), which is nearly three times the size that used in the retinal recordings. The increase in size led to an increase in the percentage of NOT neurons with a discernible WN-RF to 57.5%. In contrast, 94.7% of NOT neurons had a discernible RMO-RF. The same WN square size was used for recordings in SC and V1.

Recovering RFs from each recorded brain structure revealed that RFs recovered by using RMO were typically spatially compact and well localized, whereas WN frequently failed to produce a defined RF structure altogether (Fig. 6a–e, II; Extended Data Fig. 8). Population-level quantification confirmed not only that RMO/RCASE yielded discernible RFs in a larger fraction of neurons than WN/STA (Fig. 6f,g; Extended Data Fig. 8c) but also RFs of higher quality (Extended Data Fig. 8d). Moreover, discernible RFs emerged more rapidly with RMO/RCASE than with WN/STA across all recorded structures. This difference was particularly pronounced in NOT among the recorded brain structures, where high-quality RFs were recovered within 10 minutes of stimulation (Extended Data Fig. 8e).

This advantage was especially evident when we focused on direction-selective populations (Fig. 6g,h). For example, ∼63% of all retinal DS-RGCs had a discernible RF only with RMO/RCASE, whereas only ∼1% showed an RF exclusively with WN/STA. A similar pattern was observed for DS cells in NOT and SC, where ∼43% and ∼33% of RFs were identified exclusively with RMO/RCASE, but only ∼1% and 0% for WN/STA (Fig. 6g). Thus, despite differences in anatomical location and response properties, motion-selective neurons along the early visual system could be better identified and characterized by using the RMO stimulus.

Relative to RFs in the retina, RFs in NOT, SC, and V1 estimated using the RMO/RCASE were larger with NOT neurons exhibiting the largest RFs (Fig. 6i). This increase in RF size along the visual hierarchy has been reported and is consistent with increasing integration across visual space (Van Essen et al., 1981). In mice, previous studies have reported larger RFs in the SC and V1 compared to RGCs, and neurons of the accessory optic system were found to integrate motion signals over broad regions of visual space (Dhande et al., 2013; Niell & Stryker, 2008; L. Wang et al., 2010). Nevertheless, RMO/RCASE recovered localized RF structure in a large fraction of neurons, including populations for which WN/STA frequently failed to yield a discernible RF.

### RMO/RCASE and WN/STA are both effective in primate retina

We recorded spiking activity of 157 RGCs in two *ex-vivo* primate retinal preparations (one marmoset, one macaque retina). Similar to the mouse retina, RMO elicited higher average firing compared to WN (Fig. 6e), although the difference was less pronounced (Fig. 6a). After just 10 min of stimulation, 91.7% of primate RGCs had a discernible RMO-RF, 10.2% more than WN-RF (Extended Data Fig. 9c). Although RMO/RCASE yielded discernible RFs in a larger fraction of cells and slightly higher RF quality indices than WN/STA, the difference between the two methods was substantially smaller than in mouse retina and downstream visual structures (Extended Data Fig. 9c,d). This finding was confirmed when the two primate species were analyzed separately. In the marmoset retina, 89.8% (53/59) of RGCs had discernible RMO-RFs, vs 79.7% (47/59) for WN-RFs, whereas in the macaque retina 99.0% (97/98) and 84.7% (83/98) of RGCs exhibited RMO- and WN-RFs, respectively (Extended Data Fig. 9c). Similarly, RF quality was higher with RMO/RCASE than with WN/STA in both species, although the difference remained modest (Extended Data Fig. 9d). Moreover, with increasing stimulus duration, RF quality increased similarly with RMO/RCASE and WN/STA in both species (Extended Data Fig. 9e). This was in stark contrast to mouse RGCs, DS-RGCs, and neurons from the NOT region, where cells generally required substantially longer exposure to WN stimulation to yield a WN-RF, or where WN-RF detection failed.

These results indicate that the majority of primate RGCs, unlike mouse RGCs, were well characterized by WN analysis. We, therefore, asked whether WN-based linear models also captured primate RGC responses more effectively than they captured mouse RGC responses. We fitted linear-nonlinear (LN) models to a part of the recorded RGC response and evaluated their predictions on held-out data. We used the WN stimulus matrix as input for the LN-models trained to predict WN responses (WN-models; Fig. 6j, bottom). In contrast, we used the RCASE stimulus matrix as input for the LN-models trained to predict RMO responses (RMO-models; Fig. 6j, top). Prediction performance was quantified as the squared Pearson correlation coefficient between observed and predicted responses (ρ^2^). Across the mouse retinal population (n = 1,346 cells, 4 retinae), RMO-models achieved significantly higher prediction performance than WN-models (Fig. 6k, left). The prediction performance of the RMO- and WN-models was only weakly correlated (Fig. 6k, right; *r* = 0.19, *p* = 3.98 ⋅ 10^−12^), indicating that high prediction performance with one stimulus representation did not necessarily correspond to high performance with the other. In contrast, in primate RGCs, where RMO-models outperformed WN-models as well (Fig. 6l, left), the correlation of the prediction performance of the two model classes was stronger (Fig. 6l, right, *r* = 0.62, *p* = 2.54 ⋅ 10^−18^). These findings suggest that - considering the main primate RGC types and the stimuli employed here - both models capture largely overlapping aspects of the visual response, namely stimulus contrast.

## Discussion

Functional specialization is a feature of sensory neural systems, where different neural populations respond selectively to unique spatiotemporal stimulus patterns. Understanding how functional specialization arises along the pathways, and which neural computations are involved is a formidable experimental question. Since early processing stages respond to local unstructured stimuli and later stages to complex, structured stimuli, experimental approaches traditionally employ different stimuli to characterize the different stages. This approach renders comparisons of neural encoding between different stages challenging and hinders the characterization of neural transfer functions from one stage to the next. Here, we used a single stimulus paradigm across the early visual system and compared its efficacy in driving neural responses to that of WN.

### Structured stimuli elicit neuronal responses more effectively

In comparison to structured stimuli, WN stimuli offer the theoretical advantage of allowing for an unbiased estimation of a neuron’s response function. However, this advantage is severely compromised if the neurons do not spike in response to them. Across the systems we investigated, WN stimuli elicited the weakest response compared to a range of other stimuli. This result is surprising given that WN contains the largest contrast change per pixel and time. If the assumptions underlying WN-based system identification were met, such as linear encoding of pixel contrast, LNP models should accurately predict neuronal response behaviors. However, the LNP models we employed to capture RGC spiking responses predicted that the strongest responses would occur upon exposure to WN stimuli, which was in stark contrast to our measurements. Typically, experimental time in systems neuroscience is severely limited, and WN-RF estimates rely directly on the number of spikes elicited by WN stimuli (Schwartz et al., 2006). Given these constraints, the use of structured stimuli, such as the RMO stimulus, could greatly improve the characterization of neural response behaviors in functionally specialized systems.

### Robust responses to the RMO stimulus across early visual areas

With each processing stage, neurons encode increasingly complex features, a principle implemented in biological (Hegdé & Essen, 2000; Pasupathy & Connor, 2002) and biologically inspired artificial neural networks (Lindsay, 2021; Mineault et al., 2021). However, investigating how functional specialization emerges along a processing hierarchy presents substantial challenges. The mouse retina alone contains more than 30 retinal ganglion cell types (Baden et al., 2016; Goetz et al., 2022), each of which constitutes a distinct output channel, and many of which exhibit poorly understood response properties. Because downstream neurons often integrate inputs from multiple channels, the computations associated with a given processing stage can be difficult to isolate.

We, therefore, focused on two relatively simple retinofugal pathways, namely the retina–NOT and retina–SC pathways, which are involved in the generation of innate visually guided behaviors (Liu et al., 2016; Yilmaz & Meister, 2013), and we distinguished between non-DS and DS cell populations. We then compared neuronal responses to WN stimulation and to the RMO stimulus across these early visual stages. Across all structures, including non-DS cells in V1, neurons responded robustly to the RMO stimulus, indicating that the stimulus captures visual features that remain behaviorally relevant throughout the early visual system, while the WN stimulus was often ineffective. Consistently, RF estimation did not require a large number of spikes and could effectively be performed within 20 minutes of visual stimulation at all stages.

### Other approaches to neural system identification

WN-based reverse correlation provides an unbiased estimate of the linear component of a neuron’s response function, provided the stimulus approximates a Gaussian ensemble, and sufficient numbers of spikes are collected (Chichilnisky, 2001; Schwartz et al., 2006; Wiener, 1958). The requirement for Gaussian stimulation can be relaxed, and responses can be increased by using non-Gaussian WN variants, including binary WN (Clay Reid & Alonso, 1995), sparse noise (Komban et al., 2014), shifted noise (Pamplona et al., 2022), correlated noise (Matteucci et al., 2019), or randomly flashing bars (J. P. Jones & Palmer, 1987) and gratings (D. L. Ringach et al., 1997). A complementary line of development has been the use of specifically designed noise stimuli that feature a spatial structure that is more effective in exciting visual neurons, while still providing sufficient statistical control for system identification. The “cloud” stimulus, introduced for mouse primary visual cortex (a contrast-modulated spatially filtered noise movie with 1/f-like statistics) and later used with multiple visual cortices in the rat, evokes stronger and more population-wide responses than sparse noise alone (Matteucci et al., 2019; Niell & Stryker, 2008). Coupling cloud stimuli with multi-subunit nonlinear input models (NIM) has been shown to yield detailed characteristics of excitatory and suppressive RFs in mouse retinal ganglion cells, including ON-OFF cells, the symmetric nonlinearity of which makes them invisible to standard STA analysis (Shi et al., 2019). The common theme across these approaches is that stimulus design and analysis must be jointly optimized for the specific neural population under study. However, these methods rely on reverse correlation with the pixel contrast of the stimulus: the RF is described as a function of pixel space, and the estimated response function is the first-order (linear) component of the neuron’s computation.

It has long been recognized that stimuli beyond WN are necessary to characterize neural responses at cortical stages, and that reverse correlation can be performed against structured stimuli including natural image sequences. One example is the estimation of the linear RF of macaque V1 neurons from their responses to natural movie clips. This approach additionally revealed nonlinear boundary-selective signals that were invisible with gratings (D. L. Ringach et al., 2002). Methods for estimating spatiotemporal response functions from complex stimuli with strong multi-point correlations, such as those occurring in natural scenes and acoustic signals, were developed in both the visual and auditory domains (Klein et al., 2000; Theunissen et al., 2001; Wu et al., 2006). Information-theoretic methods (maximally informative dimensions) allow for unbiased RF estimation from responses to natural stimuli even without a Gaussian stimulus assumption (T. Sharpee et al., 2004), and cat V1 RFs adaptively change their spatial-frequency tuning depending on the input ensemble, an effect invisible with white noise analysis methods (T. O. Sharpee et al., 2006). In cat V1, natural scene animation driven by simulated eye movements elicits sparse, temporally precise spiking responses with high trial-to-trial reliability, contrasting sharply with the high variability and noise of responses to gratings or dense noise (Baudot et al., 2013). These results lead to the conclusion that structured stimuli with appropriate spatiotemporal statistics can elicit more informative and reliable neural responses than unstructured noise, even at early processing stages, such as the primary visual cortex. A popular approach for characterizing visual neurons has moved toward explicitly nonlinear statistical models that dispense with WN assumptions entirely. Generalized linear models (GLMs) fit a spiking nonlinearity directly to the stimulus-response relationship and can predict responses to arbitrary stimuli (Parker et al., 2022; Pillow et al., 2005, 2008). Multi-stage cascade models (LNLN, LNLNP) add hidden subunits that capture nonlinear spatial pooling and divisive normalization, yielding mechanistic descriptions of ON-OFF cells, direction-selective cells, and other cell types in the mouse retina (Maheswaranathan et al., 2018; Tanaka et al., 2019). At the extreme end of expressiveness, deep convolutional neural networks, trained end-to-end with neural responses to natural scenes, can match or exceed simpler models in terms of predictive accuracy for retinal and V1 neurons (Idrees et al., 2024; Maheswaranathan et al., 2023). These data-driven approaches are reviewed in (Butts, 2019). Crucially, such models naturally accommodate structured and naturalistic stimuli, which likely contributes to their improved predictive performance relative to simpler LN approaches on cells whose computation goes beyond linear filtering. However, these approaches require large datasets, extensive model fitting, and substantial computational resources, often making them impracticable for rapid characterization of neuronal populations during physiological experiments.

Within this landscape, RCASE represents what could be called a “linearized model” (Wu et al., 2006): a method in which the known nonlinear response properties of a neuron are explicitly represented in the stimulus parameterization, so that a linear model can be applied to the resulting (non-pixel) representation. Early approaches in the visual domain estimated spatiotemporal RFs of simple cells in cat striate cortex by reverse correlating responses against sequences of randomly positioned bars, which can be seen as a low-dimensional, parameterized stimulus in which “bar position” rather than “pixel contrast” was the regressor (DeAngelis et al., 1993). RCASE formalizes and generalizes this logic: the stimulus is described in terms of the presence and parameters of stimulus elements (here, moving objects), yielding a design matrix in the stimulus-element space rather than pixel space. This approach enables efficient linear estimation even when pixel-level correlations would make ordinary STA ill-posed.

A conceptually similar approach is taken by subspace reverse correlation and related feature-space methods, which improve estimation efficiency by projecting visual input onto a restricted set of predefined basis functions or stimulus features and correlating neural responses within this reduced representation, rather than estimating sensitivity independently across the full pixel space (D. L. Ringach et al., 1997; Bonin et al., 2011; Skriabine et al., 2026).

A related multi-filter extension of spike-triggered analysis, *i*.*e*., spike-triggered covariance (STC), can recover multiple linear subspaces from white-noise data, thereby capturing the symmetric nonlinearities of ON-OFF cells and complex cells that STA misses (Cantrell et al., 2010; Rust et al., 2005; Schwartz et al., 2006). STC has been used to reveal both excitatory and suppressive filter subspaces in macaque V1 complex cells and direction-selective cells (Rust et al., 2005), and has been applied to mouse retinal ganglion cells to recover ON-OFF RF structure (Cantrell et al., 2010). However, STC requires a large number of spikes (on the order of millions for full-resolution WN stimuli), making it experimentally impracticable for many recording conditions. Our STC analysis on WN data showed that: (a) STC correctly recovers the ON-OFF RF structure of a modeled LNLP cell from white-noise responses; but (b) in practice, STC fails on our experimental data because each neuron’s low WN-evoked spike count falls short of the requirements estimated from the covariance matrix dimensionality (Schwartz et al., 2006) (Extended Data Fig.5). After dimensionality reduction using the RCASE-estimated RF center to restrict the STA/STC analysis to a local spatial window (3 × 3 pixels × 10 time lags; cf. (Ahn et al., 2020; Cantrell et al., 2010)), we find that approximately 18% (WN100) and 44% (WN200) of cells show an RF recovered by the leading or trailing significant STC eigenvector (RF QI > 0.3), compared to 98% recovered by RCASE in the same cohort (Extended Data Fig. 5). These results illustrate a key complementarity: whenever the spike count permits, STC can recover multi-filter nonlinear structure from WN data, but the same data constraints that limit STA in functionally specialized neurons equally limit STC, whereas RCASE achieves high RF identification rates with substantially fewer spikes.

Constructing stimuli from nonlinear features rather than pixels, while retaining compatibility with reverse-correlation analysis, is not unique to the present approach. In the auditory system, TORC stimuli exploit the spectrotemporal structure of the auditory scene as the parameterization space for reverse correlation (Klein et al., 2000). A study of face-selective neurons used a deep generative network and genetic algorithm to evolve images that maximally elicit neurons in macaque IT cortex (Ponce et al., 2019), effectively performing optimization in a deep feature space rather than in pixel space. These examples instantiate the general principle articulated in (Wu et al., 2006): when the relevant computational dimensions of a neural population can be hypothesized or estimated, parameterizing the stimulus in that low-dimensional feature space dramatically improves estimation efficiency. RCASE applies this principle to the early visual system, where the relevant dimensions (spatial location, direction of motion, object size) are well-defined and experimentally controllable.

### Suitable stimuli for RF estimation with RCASE and their interpretation

RCASE is designed for efficient characterization of a hypothesized low-dimensional feature space. It cannot discover arbitrary unknown features because the dimensions to be characterized must be instantiated in the stimulus and encoded in the element representation. Methods that learn or adapt the feature space may therefore be preferable when the relevant dimensions are unknown. Candidate stimuli for RCASE should engage the population, sample element parameters broadly across the display, admit an element-based representation, and produce a sufficiently well-conditioned design matrix.

### Temporal correlations in the RMO design matrix

The RMO stimulus introduces temporal correlations through object motion, since each object persists across multiple frames along a deterministic trajectory. This is not the case for WN, where each pixel is independently resampled at each time step. Object motion introduces temporal correlations because each object persists across consecutive frames. Although the RCASE design matrix was approximately free of instantaneous spatial correlations at a fixed lag, kernels at different lags were not independent. The temporal RCASE kernel therefore reflects both neural response dynamics and the temporal statistics of the object trajectories. This limitation must be considered when interpreting kernel timing and apparent spatial extent. This is analogous to the situation with other structured stimuli used in retinal physiology, including moving grating sequences, where temporal correlations are similarly inherent to the stimulus design and must be accounted for in the interpretation of the results.

Future implementations could incorporate temporal whitening or trajectory-shuffling controls, although these modifications may increase data requirements or alter neural firing rates. One approach to assessing whether temporal correlations distort RF estimates is a temporal shuffling control: the frame sequence of the RMO stimulus can be randomly reordered, breaking the object-trajectory correlations while preserving the instantaneous statistical properties of each frame. Neural responses to this scrambled sequence provide a null distribution against which the structured-stimulus responses can be compared, and differences in estimated RF quality between scrambled and unscrambled conditions isolate the contribution of temporal correlations (Manookin et al., 2018). Furthermore, explicit deconvolution of the temporal correlation structure could be an approach to address this issue. Future extensions of RCASE could incorporate explicit temporal whitening of the design matrix to remove the bias introduced by object trajectories, at the cost of requiring more observations to achieve adequate signal-to-noise ratio in the whitened representation.

### The RF identification advantage of RMO/RCASE is not restricted to direction-selective cells

Because RMO contains moving contrast edges, we expected it to engage direction-selective neurons effectively. The advantage of RMO/RCASE under limited experimental time was prominent in retinal, NOT, and SC DS populations. However, RMO/RCASE also increased RF yield in non-DS populations. The WN–RMO performance difference therefore cannot be explained solely by the proportion of DS cells.

### Predicting a neuron’s response to other stimuli

The ability to utilize estimated RFs as filters to predict a neuron’s response to arbitrary stimuli defined in pixel space is a significant advantage of WN-RFs over RMO-RFs. However, for WN-RFs to serve as robust predictors, they must meet strict requirements: firstly, the neuronal responses must be strong enough to estimate the RF, and secondly, the neuron’s response behavior should be linear with respect to pixel contrast, possibly followed by a scalar nonlinearity.

In contrast, RMO-RFs are not suitable for predicting responses to arbitrary stimuli due to the non-invertible, nonlinear reparameterization of the stimulus space. However, our approach allows for the identification of RFs in functionally specialized neurons with only a few spikes. Notably, when we quantified how much of a neuron’s response to a specific stimulus could be explained by the RF, we found that predictions based on RMO correlated significantly more strongly with the experimentally observed neuronal responses than those based on WN (Fig. 6j-l). This suggests that the RFs obtained with RMO more faithfully reproduce the neuron’s response properties.

## Acknowledgements

We would like to acknowledge the following individuals and organizations for their valuable contributions to our research: We acknowledge SILABE (Simian Laboratory Europe, University of Strasbourg), in particular Pierre-Henri Moreau, for their collaboration, and provision of primate tissue for our research investigations; Wibke Schwarzer and Lucas Janeschitz-Kriegl for coordinating the primate tissue donation process; Lucas Janeschitz-Kriegl, Tiago M. Rodrigues, Akos Kusnyerik for taking the time to perform the enucleation of primate eyes used in this study; Martina De Gennaro, and Cameron S. Cowan for their support and valuable advice related to the preparation of primate tissue.

## Author contributions

M.B., M.Z., R.D., and F.F. designed the study. M.Z. performed high-density microelectrode recordings in mouse and marmoset retinae. A.B. performed high-density microelectrode recordings in the macaque retina. F.B.R. performed Neuropixel recordings in the nucleus of the optic tract in anesthetized mice. M.B., R.D., and F.F. developed the methodology. M.B., F.F. analyzed experimental data. M.B. modelled neural responses. M.B., M.Z., and F.F. designed figures. F.F., F.B.R. and A.H. acquired funding. M.B. and F.F. wrote the manuscript. A.H., R.D., M.Z., F.B.R., and A.B. reviewed and edited the manuscript.

## Funding

SNSF Eccellenza Grant PCEFP3_187001 (F.F.)

SNSF Projects in Life Sciences Grant 310030_220209 (F.F.)

SNSF Spark Grant CRSK-3_220987 (F.F.)

SNSF Sinergia Grant CRSII5_173728 (A.H.)

SNSF Sinergia CRSII5_216632 (A.H.)

SNSF Spark Grant CRSK-3_221257 (F.B.R.)

European Commission ERC Advanced Grant 694829 (“neuroXscales”, A.H.)

## Materials and Methods

### HD-MEA recordings

#### Tissue collection

##### Mice

Wild-type, male and female C57BL/6J mice aged between 8 and 30 weeks of either sex were obtained from Charles River Laboratories (Wilmington, Massachusetts, United States). They were kept under 12h light/dark cycles and group-housed in a pathogen-free environment. They were dark-adapted for 15 - 30 minutes before the experiment at which point an overdose of an isoflurane followed by decapitation was used to euthanize the animals as approved by the Basel-City Veterinary Office in accordance with the Swiss Federal laws on animal welfare. Tissue preparation was performed under dim red illumination and in an oxycarbon carbogen gas (95% O2, 5% CO2; PanGas AG, Dagmersellen, Switzerland) buffered Ringer’s Medium (110 mM NaCl, 2.5 mM KCl, 1 mM CaCl_2_, 1.6 mM MgCl_2_, 10 mM D-Glucose, 22 mM NaHCO_3_). The eyes were opened, and anterior parts were carefully removed.

##### Primate tissue

A healthy 26-month-old, male common marmoset (*Callithrix jacchus*) and a healthy 48-months-old female cynomolgus macaque (*Macaca fascicularis*) were kept and monitored in accordance with procedures approved by the French veterinary office (reference numbers; 2016112416023911 and 2021072914362019, respectively) at the non-human Simian Laboratory Europe - Silabe (University of Strasbourg, Strasbourg, France). The retinal tissue from the macaque, was kindly provided by our collaborators who euthanized the animal for a separate, unrelated ophthalmic project, which did involve precise subretinal injections to predefined parts of the retina. Healthy, non-injected retinal parts were used for recording in this study.

Enucleations of the eyes were performed by the surgeon under the supervision of the responsible veterinarian, while the blood circulation and ventilation of the animal were still in place. In short, animals were first sedated (Ketamine 1000, 10 mg/kg, intramuscularly), transported to the operating room where animals received an intravenous injection of Propofol (Propovet, 5-10 mg/kg), were then intubated and administered isoflurane gas anesthesia (ISOVET, 1-2.5%, inhalation) alongside an analgesic, Morphine (MORPHINE AGUETTANT, 2 mg/kg, intramuscularly). For the enucleation a local anesthesia (PRO-CAMIDOR, 17.3 mg/ml, 0.1 ml/eye, subcutaneously) was delivered around the orbital area. After enucleation, the animals were euthanized using a lethal dose of Pentobarbital (DOLETHAL; 180 mg/kg, intravenously).

Enucleated eyes were immediately opened, anterior parts and the vitreous body were removed, and the eye was flattened using butterfly cuts. The tissue was placed into Ames medium (#A1420; Sigma, Merck KGaA, Darmstadt, Germany), saturated with carbogen gas. The time elapsed since eye enucleation and the tissue being placed in the Ames medium was less than 5 minutes. Transport time of the samples to our laboratory was less than one hour, where a final part of the dissection (see *Tissue preparation* section for details) as well as HD-MEA recordings were conducted.

#### Tissue preparation

The retina was carefully separated from the retinal pigment epithelium and placed ganglion-cell-side down on a HD-MEA. Using a micromanipulator, the retina was pressed onto the electrodes with a transparent polyester tissue-culture membrane (10 μm thick, 0.4 μm pore size, product number 3450, Corning, New York, United States) to increase the tissue attachment to the HD-MEA and to keep the tissue in place during the recording. The membrane was modified to increase tissue perfusion with holes at regular spacing (100 μm radius in a 400 μm raster, IMPEX Leiterplatten GmbH, Michael im Lungau, Austria). The temperature was kept at 32°C with the help of an active perfusion of 4 ml/min or 6 ml/min for primate tissue and an inflow temperature controller. In case of the primate retina, a peripheral part of the tissue was used.

#### High-density microelectrode array recordings

HD-MEA recordings were performed on a CMOS-based chip consisting of 26’400 electrodes with a center-to-center pitch of 17.5 μm, and 1024 parallel readout channels(Müller et al., 2015). Each readout channel featured an analog-to-digital converter (ADC) with 10-bit precision and 20 kHz frame rate. For each retina piece, placed on the array, we first mapped the activity across the entire array to locate areas best suited for the experiment. We then selected a high-density electrode configuration in the selected region according to design constraints of the switch-matrix. Finally, the light stimulus was centered onto the middle of the recording area covering all selected electrodes. We were able to record light-evoked spikes over several hours (stable response patterns for maximum of 6 h) per retinal piece. Depending on the duration of the light stimulation protocol, the size of the active area, and real-time observations of the spiking activity, we selected suitable electrode configurations and repeated the light stimulation protocol. In other cases, we would switch to a new piece of retina from the same preparation that was kept in the dark at room temperature. We excluded all experiments, where population-wide spiking rates and patterns changed strongly over the duration of an experiment.

#### Pharmacological studies

Drug solutions including strychnine (1 μM, ID: S0532, Merck KGaA, Darmstadt, Germany), picrotoxin (100 μM, ID: P1675, Merck KGaA, Darmstadt, Germany), and gabazine (5 μM, ID: 5.05986, Merck KGaA, Darmstadt, Germany) were freshly prepared from aliquoted stock solutions stored at - 20°C. After control recordings had been obtained as described before, perfusion was switched to medium with added pharmacological agents, and a wash-in for a minimum of 20 minutes was performed. Upon finishing HD-MEA recordings with pharmacological agents a wash-out of 20 minutes was performed before starting further HD-MEA recordings.

#### Light stimulation

For light stimulation we used (1) a DLP LIGHTCRAFTER™ E4500MKII™ projector (EKB Technologies Ltd, Bat Yam, Israel), with UV (385 nm), blue (460 nm) & green (520 nm) LEDs, for experiments with mouse retinas, and (2) a DLP DLP4750LC projector (EKB Technologies Ltd, Bat Yam, Israel), with red (625 nm), green (545 nm), and blue (455 nm) LEDs for experiments with primate retina. The setup consisted of an upright microscope (Olympus BX51WI, Olympus Corporation, Tokyo, Japan) with a modified imaging path, through which we projected the light stimulus directly onto the photoreceptor layer of the retina, while simultaneously recording RGC light-evoked activity with the HD-MEA (the arrangement of the setup was described in (I. L. Jones et al., 2015)). The projected stimulus was rendered at (1) 1280 x 800 pixels resolution and a refresh rate of 60 Hz and had a final size of 3366 x 2104 μm^2^ on the photoreceptor layer or (2) 1920 x 1080 pixels resolution and a final size of 3300 x 1860 μm^2^. We adjusted for the gamma correction of the projector such that a linear relationship between light intensity and numeric color values in software was preserved. The presented stimulus was provided by (1) UV and green projector LEDs, and individual intensities of colors were first equalized with the help of computer software. Color ‘white’ thus describes equalized power of both green, presented at 10% of its full power, and UV LEDs (with a final power corresponding to 2.8x10^4^ (M-cones) and 3.1x10^4^ (UV-cones) photoisomerizations per rod per second, R*/cone/second) and no contribution from the blue LED. In primates (2), the stimulation included all red, green, and blue LEDs (corresponding to R*/cone/second of 1.7x10^4^, 1.8x10^4^ & 1.1x10^4^ for L-cones, M-cones, and S-cones, respectively) without power equilibration between them. Projected light power was measured and calculated as previously described (Farrow et al., 2013). All light stimuli were generated with Psychtoolbox-3(Brainard, 1997; Kleiner et al., 2007; Pelli, 1997) on MatLab® (v2019a or newer; MathWorks, Natick, Massachusetts, United States).

### Neuropixel recordings

#### Animal preparation

Mouse experiments were performed in accordance with standard ethical guidelines (European Communities Guidelines on the Care and Use of Laboratory Animals, 86/609/EEC) and were approved by the Veterinary Department of the Canton of Basel-Stadt. All experiments were performed on adult Hoxd10 mice (Dhande et al., 2013), male and female, with age ranging from 2 to 7 months. Mice were maintained on a normal 12-hour light/dark cycle, and group-housed in a pathogen-free environment with ad libitum access to food and drinking water. Animals were anaesthetized using a SC injection of Fentanyl-Medetomidine-Midazolam (FMM) (Fentanyl (Janssen, 0.05 mg/kg), Medetomidine (Virbac AG, 0.5 mg/kg), Midazolam (Sintetica, 5 mg/kg) in saline solution (0.9%)) and placed on a heating pad to prevent anesthesia-induced hypothermia. Eye lubricant was applied throughout the procedure to prevent corneal dehydration. The scalp was shaved using hair clippers and disinfected with Iodine solution. Local anesthesia (Lidocaine, 0.1-0.2%) was administered SC prior to incision. A ∼5 mm scalp section was removed via micro scissors. Saline solution and hydrogen peroxide were used to remove the periosteum and any remaining tissue. The scalp edges were fixed to the skull using surgical glue (Histoacryl, Braun). The skull was then etched using a micro curette to ensure cement grip. Finally, a custom-built aluminum headpost was affixed to the skull using super-glue (Loctite 403, Conrad) and dental cement (Super Bond C&B, MPE Medical GmbH). Craniotomy coordinates were marked on the skull via a permanent marker and the exposed skull covered with a silicone sealant (Kwik-Cast, World Precision Instrument). Buprenorphine (Temgesic, 0.05-0.1 mg/kg) was administered with the wake solution (Atipamezol (Virbac, 2.5 mg/kg), Flumazenil (Sintetica, 0.5 mg/kg) and administered at the end of the procedure. Analgesia treatment was continued with Buprenorphine (0.1 mg/Kg BW), administered SC every 6 hrs during light phase for three days after craniotomy, for two days after head-post mounting. During the dark phase Buprenorphine (1 ml of 0.3 mg/kg + 32 ml water) was administered in the drinking water. For the three-day postoperative period the mice were single housed. After this recovery period, mice were placed back with the littermates. On the day of first recording, the animal was anaesthetized as described above. The headpost was secured on a headpost holder and a ∼3x3mm craniotomy performed with a .7 mm burr dental drill (AEU-25 and AHP-64, Altmann Dental) above NOT, SC, or V1 location, around the stereotaxic coordinates AP -2.4-2.6 mm and ML 1.3-1.4 mm for NOT, AP -2.9-3 mm and ML 1.3-1.4 mm for SC, and -3.1-3.2, and ML 2.5-2.6 for V1, respectively. The animal was then transferred to the experimental rig, where it was placed on a heating pad to keep normothermia (37°-38°C) and its head fixed onto the headpost holder. Oxygen was administered throughout the session. Prior to probe insertion, the dura was removed and a drop of low melting point agarose (1.2%) was placed onto the exposed brain to avoid dehydration and for mechanical stability. Toe-pinch reflexes, respiratory rate >150 breath/min, temperature and heart rate were monitored throughout the procedure at 30 minutes intervals. An additional dose of FMM (25% of the initial dose) was administered after 40 minutes from induction, and in case the respiratory rate exceeded 150 breath/min, as this parameter was established to be the most accurate indicator of anesthesia level (even in absence of pedal reflex).

#### Data acquisition

The experimental rig consisted of an elevated 20 x 10 cm Thorlabs breadboard platform with a heating pad (Harvard Apparatus) on top, placed onto an anti-vibration table and facing two conjoint LCD monitors at 2560x1440 resolution. The monitors were placed in a vertical orientation, and each spanned 96x126 degrees of visual angle. The platform was clamped on a 37-cm-high pillar post, above which a probe micromanipulator (uMp-4, Sensapex) was mounted. During the experiment, the anesthetized mouse was placed on the heating pad and its head secured to the headpost holder. The mouse was positioned so that left and right eye were at a 14 cm distance from the left and right monitors, respectively, and pointing approximately at the monitors’ center. A stereomicroscope was placed to the side of the right monitor to allow monitoring of probe insertion.

Neural recordings were performed extracellularly via Neuropixel probes (Steinmetz et al., 2021) . Each probe features a 70 µm x 10 mm single shank, and 384 channels configured for recording, spanning a recording area of 3.8 mm along the dorso-ventral (DV) axis. Before the recording, a silver wire was permanently soldered to the probe ground. During the recording, the silver wire was taped onto the headpost holder, and an internal reference (the probe tip) configuration was used. The probe was mounted on a dovetail probe holder, secured to the micromanipulator, and inserted into the brain at a speed of 2 μm/s until the target depth was reached, typically around 4-4.2 mm for NOT and SC recordings, 2.5 mm for V1 recordings. For NOT recordings, stereotactic coordinates of insertion were in the range -2.4-2.8 mm on the antero-posterior (AP) axis and 1.1-1.4 mm on the medio-lateral (ML) axis. For SC recordings, coordinates were -2.9-3 mm on the AP axis and 1.1-1.4 mm on the ML axis. For V1 recordings, coordinates were 3.1-3.2 mm on the AP axis and 2.5-2.6 mm on the ML axis. The probe was left to settle for 15 minutes before the recording was started to minimize mechanical drift. Neural signals were band-pass filtered (300 – 5000 Hz) and acquired at 30 kHz via the Openephys GUI (https://open-ephys.org/) at a gain setting of 300x.

Stimuli were presented on two gamma-corrected monitors with a refresh rate of 60 Hz. NOT was localized online by the presence of robust direction-selective responses of the recorded neurons to moving gratings (spatial frequency, 0.05 and 0.1 cycle-per-degree; speed, 10 and 45 °/s) in the nasal direction. V1 and SC were localized online via the cells’ retinotopic location, based on (Garrett et al., 2014; Mrsic-Flogel et al., 2005) in response to a moving bar (20° size and 45 °/s speed) at different locations eventually covering the whole monitor.

#### Conversion of visual angle to distance on the retina

The visual angle (θ) was determined using the formula θ = 2 *arctan(S/2D), where *s* represented the size of the stimulus (measured in degrees) and *D* the distance between the mouse eye and the screen (14 cm). The visual angle was then converted into *µm* on the retina based on an approximated eye size of 3300*µm* (Remtulla & Hallett, 1985) as reported previously (Schmucker & Schaeffel, 2004).

#### Histology and immunohistochemistry

At the end of the recording session, the probe was extracted, coated with a fluorescent dye (DiI, Thermo Fisher, V22885) and reinserted at recording depth. Then, mice were placed under overdosing Isoflurane anesthesia and underwent transcardiac perfusion with PBS followed by 4% paraformaldehyde (PFA). The brain was then extracted and kept in PFA for 24 hours, after which it was moved to phosphate buffer solution (PBS) overnight. The brain was sliced with a vibratome, and the slices (75 µm thickness) were placed in PBS.

PBS was then removed, and samples were incubated in 300 μl blocking solution composed of 10% normal donkey serum (NDS; Sigma-Aldrich, S30-M), 1% BSA (Sigma-Aldrich), 0.02% Sodium Azide (NaN_3_; Sigma-Aldrich, S2002), 0.5% Triton X-100 (Sigma-Aldrich, 93443) and 1× PBS for 2 h under shaking conditions at room temperature. For antibody incubations, the same buffer was used with 3% NDS. Samples were then incubated for 2 days at room temperature under shaking conditions in primary antibody solution (1:200 GFP anti-rabbit polyclonal, *Thermofisher*). After three PBS washes, the secondary antibody solution (1:200 Alexa-488 conjugated donkey anti-rabbit IgGV, *Thermofisher*) was applied and left overnight under shaking conditions at room temperature. The following morning, the slices were washed with PBS, placed on glass slides, and mounted with ProLong (*Invitrogen*). Images of the brain slices were acquired using a Spinning Disc Confocal system (*Yokogawa Electric*) on a microscope operated by the *CellSens* Software (*Olympus*). The NOT was identified by the presence of GFP-fluorescent axonal terminals below the cortical layers and anterior to the Superior Colliculus.

### Light Stimuli

#### Light stimulus and retinal location

Light-evoked RGC spiking acitivity was recorded from randomly sampled portions of murine retinal preparations without systematic selection for dorsal or ventral positions. The equalized UV–green stimulation was designed to activate both the UV-dominant dorsal and green-dominant ventral retinal fields. Because the composition of RGC types varies across the dorso-ventral axis, we refrained from making claims about the absolute population proportions of RGC types in our recordings.

#### Random moving objects (RMO) stimulus

The newly developed light stimulus consisted of moving square-shaped objects. Each moving object was given a set of parameters determining its appearance and movement trajectory. The parameters of the individual objects were randomly selected from a uniform distribution across the preselected stimulus parameter space, which could be defined on a continuous range (direction of movement in range [0-360°]) or on a predefined set (e.g., 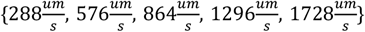). For example, the trajectory of an object would be determined by a random starting location on the screen (x, y) and in time (t), and its direction and speed of movement. Its appearance would be determined by size, and/or color parameter. To generate uniform motion trajectories in the RMO stimulus, we determined the maximal distance that an object could travel given the selected speed parameters during its lifetime (3 seconds). We then randomly selected starting points of object trajectories uniformly across an area which was larger than the area covered by the screen. This procedure ensured that the same number of objects entered and left the screen, resulting in uniform object density across the entire screen area. In initial experiments the density of objects, i.e. the number of simultaneously present objects in a single frame was varied. For most results reported here, an average of roughly 7 objects was present at the same time, resulting in an average coverage of roughly 4% for objects with a size of 200 µm edge length. Depending on the question we chose different parameterizations for the RMO stimulus (see Extended Data Table 2).

#### Handling of object occlusions

When generating the RMO stimulus, objects were drawn into each stimulus frame sequentially and overwrote the pixel values at the locations at which they were visible. In experiments with a single object color, overlapping objects, therefore, did not add their contrast, and the contrast remained at its full value. In experiments with both bright and dark objects, the last object drawn occluded previously drawn objects at the overlap locations.

To quantify how frequently objects occluded one another within a cell’s RF, we approximated both the shape of each object and the shape of the RF as circles with a diameter of 200 µm, with the RF placed at the center of the display and analyzed a 20-min RMO stimulus. For each stimulus frame, an object was considered to be within a given RF if the center position of the object fell within 200 µm (i.e., the sum of the object and RF radii) of the RF center location. When multiple objects were considered within a single RF within the same frame, they were grouped into pairs (two objects within the RF at the same time), and triples (three objects within the RF at the same time). We did not consider groups of more than three objects since they were too rare. For each unique group of objects, the minimum center-to-center distance between any object of the group and the RF attained over the entire stimulus duration was retained. A pair was classified as touching if this minimum distance was below 200 µm (twice the object radius) and as half-occluded if it was below ≈ 80.8 µm (i.e., the center-to-center distance at which the overlap area of two equal circles equals half the area of a single circle). Fewer than 3% of all object pairs were ever half-occluded. Object collisions/occlusions were only considered in the neural response analysis to pairs of objects (Fig. 5); in all other analyses, occluding objects were not treated differently from isolated objects, as we verified that their presence did not measurably affect the estimated spatial RFs.

#### Random static objects (RSO) stimulus

The static random objects stimulus was generated in the same framework as the RMO. Here, the speed parameter was set to zero. On average 5.3 objects (200 µm edge length) were present in a single frame, resulting in a coverage of ∼3%. Each object was presented for 3 frames similar to the 20 Hz refresh rate of the dense WN stimuli. The obtained stimulus effectively was a sparse WN stimulus, which we analyzed in the RCASE framework. The RSO stimulus was only presented for the mouse retina.

#### Other light stimuli

In addition to the RMO and RSO stimuli, the following light stimuli were used for the mouse retina:

i. Full-field chirp stimulus, consisting of three main components, (a) a 3 second full contrast on-off step, (b) a full contrast, frequency modulation sweep, and (c) a fixed frequency, contrast modulation sweep(Baden et al., 2016).
ii. Moving bar stimulus. A bright bar spanning the entire display area with a length of 620 *µm* moving at a speed of 1296 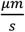 in 16 different directions; 4 trials per direction. To ensure that the total light intensity was the same for all directions, the stimulus was shown within a circular aperture of 700 μm or larger radius.
iii. A contrast step and flash stimulus consisting of the following parts: (1) 1s full-field black, (2) colored flash for one frame (1/60 s), (3) 1s full-field black, (4) 1s full-field color, (5) black flash for one frame, (6) 1s full-field color, (7) 0.5s full-field black; 4 trials per color (White/UV/Green)
iv. A binary white noise (‘random checkerboard’) stimulus with a temporal resolution of 20 Hz and square edge size of either 100 *µm* or 200 *µm* with a duration of 20 minutes. For comparing the responsivity of the retina to different combinations of checkerboard properties, we used sizes of 30, 50, 100, 200, 400 µm and temporal frequencies of 10, 20, 30, 60 Hz. For comparing the response reliability across repetitions of the same stimulus, the duration of the checkerboard was limited to 1 minute.
v. A spatially correlated ‘cloud’ stimulus as described in ref. (Shi et al., 2019). In short, a random checkerboard with 45 µm squares was filtered with a two-dimensional spatial Gaussian filter centered at the origin (in Fourier space) with a standard deviation of ∼1.3 cycles/mm. We presented the cloud stimulus in two versions: the cloud stimulus itself, and a ‘normalized cloud’ stimulus in which the filtered cloud was rescaled to the range [−1,1] to maximize its contrast.

The stimulus set for recordings in the mouse NOT consisted of:

i. Moving gratings (spatial frequency, 0.05 and 0.1 cpd; speed, 10 and 45 °/s)
ii. A full-field chirp stimulus (as described above (*iii*))
iii. The RMO stimulus (parameters see Extended Data Table 2), with the exception of the comparison of object size (Extended Data Fig. 10b, c), which was either 20° (≈576 µm) or 200 µm (≈7°).
iv. The binary white noise (pixel edge size 10°, 20Hz) with the exception of the comparison of pixel edge size (Extended Data Fig. 10b, c), which was either 10° (≈288 µm) or 100 µm (≈3.5°).

For the marmoset and macaque retina the stimulus set consisted of the RMO stimulus (parameters see Extended Data Table 2), a full-field chirp stimulus, and binary white noise (Marmoset: pixel edge size 60 *µm*, 30Hz; Macaque: pixel edge size 100 *µm*, 20 Hz).

### Spike sorting

The recorded extracellular action potentials of RGCs with the HD-MEA were spike-sorted as described in (Diggelmann et al., 2018). Briefly: The recorded electrode array area was split into local groups of nine or fewer electrodes, each of which was treated as an independent measurement. The following steps were performed for each of these groups separately and in parallel. The voltage signal from each electrode was bandpass-filtered (0.3 - 6 kHz), and spikes were detected when the signal crossed a threshold of 4.2 times the noise standard deviation. The spatiotemporal waveform of each detected spike was cut out of the signal and saved. After dimensionality reduction, an unsupervised clustering method (mean-shift clustering) was used to find characteristic spike templates for different neurons. All spikes were matched to the best-fitting template, and, finally, clusters of spikes with similar waveforms were merged together. Because an RGC could be detected on multiple electrode groups, automatic duplicate detection was employed. It used average spike waveforms and spike time similarities as a criterion to flag and remove duplicates. We used a two-step spike curation procedure, one automatic and one manual, to ensure inclusion of well-sorted units only. A unit passed the automatic curation procedure if all of the following conditions were sustained: (1) Mean spike amplitude > 8*σ* from the mean of the raw traces, where *σ* was the estimated standard deviation of the noise; (2) a Gaussian fit of the spike amplitude histogram explained at least 80% of the variance; (3) the cell responded towards each stimulus with more than 5 spikes; (4) the firing rate over the whole recording exceeded 0.1 Hz. In the second step units were manual curated using a self-developed GUI by the following criteria: (1) Amplitude histogram was Gaussian, (2) similarity of waveform shape, and (3) unit was not split or showed multi-unit activity. Refractory-period (inter-spike-interval, ISI) violations were not used as an explicit curation criterion, but ISI distributions were inspected during pipeline validation; across the RGC population the median fraction of spikes with ISI < 1.5 ms was 0.15% ± 0.3% (mean ± SD), confirming well-isolated units.

For spike sorting of Neuropixels (Steinmetz et al., 2021) data from the mouse NOT we used Kilosort2.5(Pachitariu et al., 2024; Steinmetz et al., 2021) with default parameters, followed by offline spike curation using *Phy* (https://github.com/cortex-lab/phy).

### Data Analysis

Data analysis was performed using MATLAB 2023a and Python 3.8.

### Contrast modulation per pixel

To compare the light stimuli with respect to their overall rate of contrast change (Fig. 1c,d), we computed the average contrast change per pixel and second for each stimulus. The stimulus movie was represented as 8-bit gray values *I*(*x, y, t*) ∈ [0,255]. For each pixel we computed the absolute frame-to-frame difference ∣ *I*(*x, y, t* + 1) − *I*(*x, y, t*) ∣ and normalized by the maximum, so that the largest possible contrast change of a single pixel equaled 1. We then summed over all pixels and frame transitions, divided by the number of pixels and the number of frame transitions, and multiplied by the stimulus frame rate (60 Hz) to obtain the average contrast change per pixel and second.

### Firing rate estimation

To calculate the average firing rate of a cell during a stimulus, we divided the number of spikes during the stimulus by the duration of the stimulus. We calculated firing rates as a function of time for each trial via kernel density estimation (Grün & Rotter, 2010) using a gaussian kernel and then averaging the firing rate over trials. Parameters used for the standard deviation of the Gaussian kernel *σ* and sampling rate Δ*t* varied across analysis methods and are reported in the corresponding method sections.

### Estimation of the maximum firing rate

In addition to the time-averaged firing rate, we estimated the maximum firing rate of each mouse RGC for every light stimulus (Fig. 1d). For each cell, the spike train recorded over the full duration of the stimulus was binned at a temporal resolution of 1 ms and convolved with a 100 ms boxcar window (i.e., spikes were counted in a 100 ms window slid across the spike train in steps of 1 ms). The resulting sliding spike counts were divided by the window length (100 ms) to obtain instantaneous firing rates in Hz, and the maximum over the entire stimulus was taken as the cell’s maximum firing rate. The reported value is the median of the maximum firing rates over the cell population, with its uncertainty estimated by bootstrap resampling.

### Estimation of the uncertainty of the median via bootstrapping

The uncertainty of reported median values (e.g., the median firing rates in Fig. 1d,g and Fig. 6) was estimated by bootstrap resampling. For each bootstrap iteration, we drew, with replacement, a sample of the same size as the original dataset and computed its median. This procedure was repeated 10,000 times, and the standard deviation of the 10,000 bootstrapped medians was reported as uncertainty (error bars).

### Reverse correlation against stimulus elements (RCASE)

To analyze neural responses towards the random moving objects (RMO) and static random objects (RSO) stimuli, we used reverse correlation against stimulus elements (RCASE). RCASE starts by finding a nonlinear reparameterization of the stimulus in terms of the presence or absence of stimulus elements (SE). SEs were characterized by their position on the screen and a set of parameters that described their appearance. In the RMO stimulus SEs were squares, which we parameterized by their location, size, speed, direction of movement, and color. The precise set of parameters used depended on the experiment. For the RSO stimulus, SEs were white squares, parameterized only by their location. The stimulus was defined by a set of frames at 60Hz. Each SE would appear at a random frame and then stay on the screen for several frames (3s for the RMO stimulus, 0.05s for the RSO stimulus). For the RMO stimulus, each moving object was encoded by a set of SEs. SEs belonging to the same object had the same value for all parameters except for the location on the screen. If they were in consecutive frames, they could have a different location (because the object moved) or the same location (because space was binned and the object might stay in the same spatial bin for multiple frames depending on its speed and the spatial bin size).

To construct the stimuli, the starting location of each object was chosen uniformly at the resolution of the display, i.e., uniformly over pixel coordinates. The direction of movement was chosen uniformly between 0° and 360°. We binned the location and direction parameters of SEs to construct the stimulus design matrix X. For the location, the bin size was chosen such that it matched the WN pixel size of the respective experiment (mouse retina 100*µm* or 200*µm*; mouse NOT 10° visual angle; marmoset retina 60*µm*; macaque retina 100*µm*). We binned the direction of movement, with a bin size of 22.5°, which matched the 16 directions of the moving bar stimulus. The other parameters (color, size, speed) were chosen from a discrete set during the stimulus generation and required no further binning for the analysis. To summarize, an object was defined by a set of parameter values and a set of SEs. Each SE was associated with a single stimulus frame in which it was present on the screen. SEs belonging to the same object were in consecutive stimulus frames (i.e., the object appeared once and disappeared once) and had the same parameter values for all frames, except for the location parameter.

In a single stimulus frame *i* and for each SE, we encoded the SE parameters using one-hot vectors 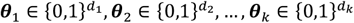 , where *d*_*k*_ was the dimensionality of the parameter space *k*. For example, a SE with the speed of 576μm/s and a speed parameter space [288μm/s, 576μm/s, 864μm/s, 1296μm/s, 1728μm/s] was encoded by the row-vector *θ*_*s*_*peed* = [0 1 0 0 0]^*T*^. For each SE present in frame *i*, we constructed a SE-tensor by the tensor product of the one-hot vectors:

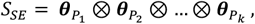

Where 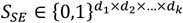.

The SE-tensors of all SEs present in frame *ii* were then summed:

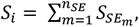

where 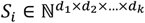 and *n_SE_* was the number of SEs present in frame *i*. The full stimulus tensor *s* was obtained by concatenation of *S_i_* over all stimulus frames T, such that 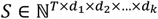. The tensor *S_i_* was vectorized into a 1-dimensional vector *xi* ∈ ℕ^*D*×1^, where *D* = ∏ *d*_*k*_ was the total stimulus dimensionality. We constructed the sparse RCASE stimulus design matrix *X* ∈ ℕ^*T*×*D*^ by concatenating the *x_i_* over frames:

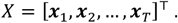

By construction, for *T*→ ∞ :

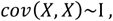

where I was the identity matrix. To calculate an RF estimate, the neural response needed to be correlated with the stimulus design matrix *X*. To this end, and identical to how we processed neural responses for the WN analysis (see below), we binned the time of the neural response with the frame rate of the stimulus, so that the neural response of a single neuron was represented by a vector of spike counts *y* with the same number of elements *T* as rows in *X*. We then calculated RFs by linear regression (reverse correlation) of the stimulus design matrix with the temporally shifted neural response *y_τ_* , where *τ* was the temporal shift or ‘lag’, defined as a multiple of stimulus frames. We performed the reverse correlation for different time lags in the range of [-2s, 2s] with 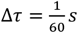:

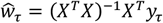

*ŵ_τ_* was the RCASE RF for a given time lag *τ*. Because the stimulus was high-dimensional but sparse, we calculated *P* = (*X*^*T*^*X*)^−1^*X*^*T*^ only once for all time lags using Matlab’s sparse matrix multiplication and then shifted the neural response for each time lag. This approach was valid since either shifting the response or the stimulus is equivalent (D. Ringach & Shapley, 2004). Because objects were present over multiple stimulus frames, the stimulus design matrix was highly correlated in time, which in turn led to stimulus correlations in the estimates of *ŵ_τ_* over different values of *τ*. We did not attempt to correct for temporal correlations of the stimulus, and the stimulus correlations must be considered when interpreting the obtained RFs. Finally, RF estimates for different time lags were concatenated to obtain the full RCASE RF *ŵ* ∈ ℝ^*H*×1^, where *H* = *D*⋅ *d_τ_*, and *d_τ_* was the number of time lags which we considered. *H* was reshaped to obtain the multi-dimensional RF tensor:

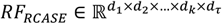

Depending on the experiment and the analysis, we reduced the parameter space of the stimulus design matrix *X* by summing over a set of parameter dimensions. This resulted in higher signal-to-noise ratios (RF quality) for the remaining parameter dimensions. To estimate spatiotemporal RFs, we summed over all parameter dimensions apart from the location parameters. To estimate tuning curves to a specific parameter *θ*, we summed over all parameter dimensions apart from the location parameters and the parameter *θ*.

### White-noise analysis

We defined the WN stimulus design matrix *X* at a 20Hz (mouse) or 30Hz (marmoset/macaque) frame rate. Each row *X_i_*,: of *X*, represented the vectorized 2-dimensional WN stimulus of frame *i*. The values of *X* were chosen uniformly at random from {-1,1}, which represented black or full contrast, respectively. By construction:

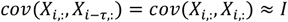

Identically to the RCASE analysis, we represented neural responses by vectors of spike counts *y* at the WN frame rate with the same number of elements *T* as rows in *X*. RFs for time lag *τ* were estimated using the spike-triggered average (Schwartz et al., 2006), which is proportional to reverse correlation of X with y, and to linear regression of X with y:

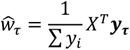

*ŵτ* was the WN RF for a given time lag *τ*, and *y_τ_* was the neural response shifted by a lag of *τ* frames. In total 15 time lags were included to compute the full WN spatial RF.

### Analysis of Cloud and normalized-cloud stimuli

RFs for the cloud and normalized-cloud stimuli were estimated in the same way as the binary white-noise stimulus with their respective stimulus design matrices.

### Decomposition of spatiotemporal RFs into spatial and temporal components

We rearranged the 3-dimensional spatiotemporal (*x, y*, *τ*) RCASE and WN RFs into two-dimensional matrices, where rows denoted the spatial dimension, and columns denoted the temporal dimension: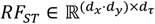. We then applied principal component analysis (Jolliffe & Cadima, 2016) (PCA) to the spatiotemporal RFs and projected the spatiotemporal RF on the first principal component. The obtained vector was then arranged in a two-dimensional matrix with the original spatial dimensions, which we defined as the spatial RF: 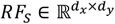. We determined the location of the RF center as the coordinates of the pixel in the spatial RF with the largest absolute value. The temporal RF or temporal kernel 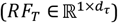 was defined as the time course of the spatiotemporal RF at the location of the RF center.

### Estimation of RCASE tuning curves

We calculated RCASE tuning curves with respect to a single SE parameter *θ*. To this end, we calculated the 2-dimensional matrix 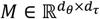, by summing the full stimulus tensor *s* over all other parameter dimensions apart from *θ* before using RCASE to estimate the parameter-dependent spatiotemporal RF. *M* represented the time courses of the neural response at the RF center for each value of the parameter *θ*. We obtained tuning curves by singular value decomposition of *M*, similar as described in (Baden et al., 2016):

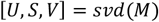

The first column of *U* was taken as the tuning curve.

### Parameter-dependent RFs and tuning curves

To illustrate how the estimated RF depends on the choice of stimulus-element parameters, we estimated parameter-dependent spatiotemporal RFs for individual stimulus-element parameters (Fig. 4). For each value of a parameter *θ*, the parameter-resolved spatiotemporal RF, its spatial and temporal components, and the tuning curve with respect to *θ* were obtained as described above. For visualization purposes, the spatial RFs for the different values of *θ* were normalized jointly across all values of *θ* for a given neuron. For this normalization, we used a single shared minimum and maximum across all values of *θ*, so that differences in RF amplitude across parameter values were preserved. For the direction parameter, the direction of movement was binned into 8 bins for this analysis.

### Peri-object time histogram analysis

The methods described so far were employed to determine the RMO RFs including the location of the RF centers. By utilizing a cell’s RF center, we attributed additional properties to the objects of the RMO stimulus, such as the distance *d*_0_ with which a moving object passed each RF center, and the time point *t*_0_ at which the object was closest to the RF center. The following procedure was conducted for each RGC individually: For each object, we defined the ‘closest approach time point’ *t*_0_ as the time point at which the object had its minimal Euclidean distance *d*_0_ in visual space to the RGC’s RF center. Subsequently, we defined distances for objects that passed to the left of the RF center as negative distances. All spike times in a time interval around *t*_0_ were shifted in time such that *t*_0_ = 0 and assigned to the object as its own ‘object spike train’. We then estimated the RGCs response profile *R*(*d*_0_, *t*) to the moving objects as a function of distance *d*_0_ and time *t* relative to *t*_0_ by binning the object spike trains in time and distance and averaging over objects. Summing *R* over a distance interval [−*d_max_, d_max_*] yielded the peri-object time histogram (POTH).

### Analysis of the neural response to pairs of objects

We removed all objects for which |*d*_0_| > 100*µm* for this analysis. For the remaining *N* objects, we sorted the closest approach time points 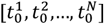, 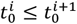 which defined the inter-object times 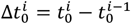, with the property, by definition, that 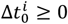. This procedure selected object pairs, consisting of object A and object B, which passed the RGCs RF center with different inter-object times 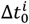 . Object A moved first over the cells RF followed by object B. We grouped the B objects of all object pairs with 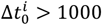 m_S_ and defined them as control objects. We binned the inter-object times into bins of 100 m_S_ (Δ*t* in Fig. 5b).

### Estimation of distance tuning curves and comparison of RF size in mice

RMO RF sizes were estimated for each cell by utilizing *RR*(*d*_0_, *t*) for *t* ∈ [−0.5 *s*, 1 *s*] with time bins of Δ*t* = 10 *ms* and *d*_0_ ∈ [−500 *µm*, 500 *µm*] with distance bins of Δ*d*_0_ = 50 *µm*. We smoothed the resulting matrix *RR*(*d*_0_, *t*) with a Gaussian kernel 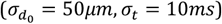 and upsampled it ten-fold using Matlab’s *imresize* function. We then calculated a distance tuning curve by singular value decomposition [*U, S, V*] =*svd*(*R*), as the first column of the *U*. The RF size was determined as twice the standard deviation *σ* of a one-dimensional Gaussian fitted to the distance tuning curve minus half the object edge length (100*µm*).

We fitted a two-dimensional Gaussian to the spatial WN RF of each cell. The WN-RF size was then defined as two times the geometrical mean of the standard deviations along the major and minor axis (Field et al., 2007) of the fitted Gaussian.

### Estimation of RF size for the comparison across visual areas

To compare RF sizes across processing stages (Fig. 6i), we estimated the RMO-RF size by fitting a two-dimensional Gaussian to the respective spatial RF. In contrast to the RMO-RF size used for the comparison between WN and RMO RF size, which was derived from the peri-object time histogram, here the RMO-RF size was obtained directly from the RMO spatial RF. The RMO-RF size was defined as two times the geometrical mean of the standard deviations along the major and minor axis of the fitted Gaussian.

### RF Quality index

We quantified the RF quality utilizing a RF quality index (Baden et al., 2016). A two-dimensional Gaussian was fitted to the spatial RF, using Matlab’s *lsqcurvefit*. The RF quality was defined as one minus the fraction of variance unexplained by the Gaussian fit 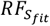:

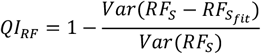

A cell exhibited a RF if the quality index exceeded a value of 0.3. The same index was computed for each stimulus and method on its respective spatial RF estimate, i.e. the spatial component of the WN STA for the WN-RF, and the spatial component of the RCASE estimate for the RMO-RF.

### ON-OFF index

We defined an ON-OFF index based on the RGCs’ responses to the full-field chirp stimulus to distinguish between ON-, OFF and ON-OFF cell types. The ON-OFF index was calculated by comparing the number of spikes in response to the first contrast step (a positive, ON, contrast step) to the number of spikes in response to the second contrast step (a negative, OFF, contrast step):

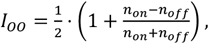

where *n_on_* was the number of spikes during the ON contrast step (3s), and *n_off_* was the number of spikes during a OFF contrast step (3s). The ON-OFF index was scaled to the interval [0,1], where 0 indicated a strong OFF-response, 0.5 indicated balanced ON and OFF responses, and 1 indicated a strong ON-response.

### Direction-selectivity index

A direction-selectivity index (DSI) was computed to determine DS-RGCs in the mouse retina and NOT. For the moving bar stimulus, firing rates were computed for each trial and 16 directions (Gaussian Kernel, Δ*t* = 0.01_*s*_, *σ* = 0.05_*s*_). Tuning curves *c*(*α*) ∈ ℝ^16×1^ as a function of direction *α* were computed by averaging over trials and summing over the time-dimension. The DSI (Baden et al., 2016) was then computed by projecting the tuning curve on a complex exponential 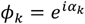, where *α*_*k*_ was the k-th direction, and normalizing it by the summed tuning curve:

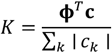

The DSI was then defined as the magnitude of the complex number *K*, yielding a value between 0 and 1:

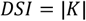

Significance of the DSI was determined by a permutation test. For each cell, direction-dependent responses were arranged in a direction-trial response matrix by summing over the time-dimension. Entries in this direction-trial matrix were randomly permuted (both over trials and over directions), destroying any relationship between the stimulus and the response, before computing the permuted DSI. We repeated the process 1,000 times to create a null distribution for the DSI. The p-value was defined as the fraction of permuted DSIs greater than or equal to the observed DSI. A DSI with *p* < 0.05 was determined as significant.

To compute the corresponding DSI for the RMO stimulus, we first calculated the stimulus element tensor 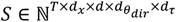 where *θ*_*dir*_ was the parameter for the direction of movement. For each cell, we determined the RF center (*x*_*RF*_ , *y*_*RF*_ ) and the temporal lag *τ*_*opt*_ at which the temporal RF was maximal. A two-dimensional stimulus design matrix for the direction parameter, 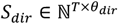, was obtained by *s*_*dir*_ (*t, θ*) = *s*_*SE*_ (*t, x* = *x*_*RF*_ , *y* = *y*_*RF*_ , *θ*, *τ* = *τ*_*opt*_ ). The direction tuning curve was computed as *c* = *s*_*dir*_ ^*T*^ *y*, where each element of the tuning curve was normalized by dividing by the number of objects for the corresponding direction. The DSI was then calculated with the same procedure as for the moving bar. Significance of the DSI was established in a similar fashion as described for the moving bar. Here, the entries of the two-dimensional direction matrix *s*_*dir*_ were shuffled 1000 times, to calculate the null distribution for the DSI.

A cell was defined as direction-selective if its DSI exceeded 0.33 and was statistically significant (p < 0.05). For the mouse retina, this criterion was required for both the moving-bar and the RMO stimulus. For the NOT, SC, and V1, the moving-grating direction-selectivity index was used.

### Estimation of preferred direction and definition of direction-selective cells

Preferred directions were estimated by fitting the direction tuning with a von Mises function,

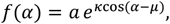

where *µ* is the preferred direction, *κ* the concentration parameter and *V* the amplitude. The preferred direction was taken as *µ*, i.e., the location of the peak of the fitted curve.

### Response-reliability analysis

We quantified the trial-to-trial response reliability of mouse RGCs to the different light stimuli (Fig. 1f, Extended Data Fig. 1,2). For these experiments, the RMO and WN stimuli were each presented as 9 repeated 1-min trials in an interleaved fashion (1 min WN followed by 1 min RMO per trial), and the moving-bar and chirp-sweep stimuli were presented with four and six trials, respectively. For each trial, time-resolved firing rates were computed by kernel density estimation as described above (Gaussian Kernel, Δ*t* = 0.1 *s*, *σ* = 0.05 *s*). Reliability was defined as the Pearson correlation coefficient between the binned single-trial firing rates across all pairs of trials: for each cell, the responses of all 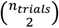 trial pairs were concatenated into two vectors, one holding the first and the other the second trial of each pair, and these two vectors were correlated (Cowan et al., 2020).

To avoid the response transient at stimulus onset (Extended Data Fig. 1b-d), the first 10 s of each 1-min RMO and WN trial were discarded, leaving 50s of stimulus; the moving bar (four trials, full epoch duration of ∼3.6 s) and chirp sweep (six trials, 32 s) were analyzed over their full duration. For the sliding-window analysis, to compare reliability across stimuli of different duration and trial number, the same number of trials (4) was used for all stimuli. The correlation coefficient was computed in 1,000 ms windows (10 bins of 100 ms) of the trial-resolved firing rate, sliding the window across the stimulus in steps of 100 ms, and the maximum correlation coefficient over all window positions was taken as the reliability value. The same reliability measures were used to compare mouse RGC responses to the RMO stimulus presented at a low and a high object density (Extended Data Fig. 3a–d): the two density variants were each shown as 10 interleaved 1-min trials, and trial-to-trial reliability was quantified both over the complete trial and with the 1,000 ms sliding-window analysis described above.

### Linear-Nonlinear Poisson Models of RGC responses

We simulated RGC responses to the stimuli employed in this study using a parameterized Linear-Nonlinear Poisson model (LNP model), consisting of a linear spatiotemporal filter, a static nonlinearity, and an inhomogeneous Poisson process for spike generation. The linear stage of the model consisted of a spatiotemporal filter, that was obtained by the tensor product of a two-dimensional spatial filter and a one-dimensional temporal filter. The spatial filter was modelled as a 2-dimensional Gaussian:

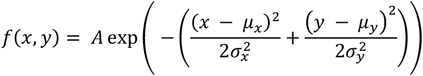

Where *x, y* denoted the location in space where the function was evaluated, *A* was the amplitude, (*µ*_*x*_, *µ*_*y*_) described the center of the Gaussian, and *σ*_*x*_, *σ*_*y*_ were the x and y standard deviations of the Gaussian. We only considered circular Gaussians, i.e., *σ*_*x*_ = *σ*_*y*_. The amplitude *A* was set to 1. The center location (*µ*_*x*_, *µ*_*y*_) were chosen uniformly over the visual space in steps of 50 *px*, avoiding an exclusion zone of 100 px at the borders. For the RF size estimation analysis, RF sizes were chosen uniformly from 25*µm* to 500*µm* in 25*µm* steps. To convert between px and *µm* the same conversion factors were used as in the HD-MEA setup. A parameterized Gabor function was chosen as the temporal filter (Dayan & Abbott, 2001; Jouty et al., 2018)

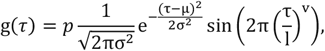

where *p* was the polarity (1 for ON vs. -1 for OFF), *v* was the “speed” determining how quickly the model responded to contrast changes, and *l* was the “length” of the temporal filter, which determined over how much time the filter integrated the stimulus. We fixed *p* = 1, *v* = 0.6105, *l* = 0.4 for all model cells, corresponding to transient ON-cells. Except for Fig. 1g (n = 110 cells per cell type), Extended Data Fig. 4g and i, where the following parameters were chosen for each cell type: transient ON-Cells *p* = 1, *v* = 0.6105, *l* = 0.4 , transient OFF-Cells *p* = −1, *v* = 0.6105, *l* = 0.4 , sustained ON-Cells *p* = 1, *v* = 1.2, *l* = 0.4. The model’s spatiotemporal RF was then obtained as follows:

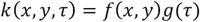

The RMO and WN stimulus were saved as videos in the resolution of the projector used in the mouse retina recordings (1280x800). For each model cell, stimulus videos were normalized between [-1,1] and cut around the RF center in a square with edge length of ∼1.25x the RF size. The cut RF videos were downsampled to a resolution of 51x51 pixels, where they exceeded this resolution, to reduce computational cost. The linear response *L* of the model was computed by convolving the spatiotemporal filter with the zero-padded stimulus movie *s*(*x, y, t*) (Dayan & Abbott, 2001) ,

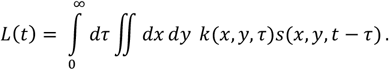

A rectified linear unit activation function was used as static nonlinearity:

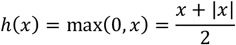

The instantaneous firing rate *y* ∈ ℝ^*Tx*1^ was computed as

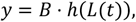

where B is the firing rate amplitude and was set to 50Hz. Poisson spike trains were generated in discrete time bins 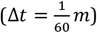 based on the instantaneous firing rate *PP*(_*s*_*pii*_*k*_*e*_*s*_|*y*) = *y*(*t*)· Δ*tt*.

### LNP model with a variable response threshold

To investigate how the threshold of the spike-generating nonlinearity affects the identifiability of a cell’s RF (Extended Data Fig. 4e,f), we simulated responses of an LNP model with a variable threshold. The model had the same architecture as described above, with the following modifications. The spatial filter was modelled as a difference of two Gaussians (DoG):

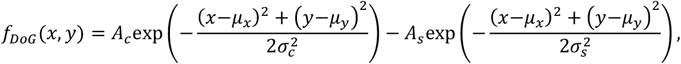

where the subscripts *c* and _*s*_ denote the center and surround Gaussian, *A*_*c*_ , *A*_*s*_ are their amplitudes and *σ*_*c*_ , *σ*_*s*_ their standard deviations. On a 10 × 10 pixel grid, both Gaussians were centered at *µ*_*x*_, *µ*_*y*_) = (5,5, with *σ*_*c*_ = 1 and *σ*_*s*_ = 3 pixels, and *A*_*c*_ = 1 and *A*_*s*_ = 0.5. The temporal filter was the Gabor function described above ( *p* = 1, *v* = 0.65, *l* = 0.4 ), evaluated at 12 time points over *τ* ∈ [0,0.5] s, yielding a 10 × 10 × 12 spatiotemporal filter. As the static nonlinearity we used a thresholded rectified linear unit, *h*(*x*) = *max*(0, *x* − *thr*) , where the threshold *thr* was varied from 0 to 0.975 in steps of 0.025. Raising the threshold shrinks the rectified response and would, on its own, drive the peak firing rate toward zero; to compensate for this issue and to keep the peak firing rate approximately constant across thresholds, the firing-rate amplitude *BB* in *y* = *B* ⋅ *h*(*L*(*t*)) was made threshold-dependent, *B* = 100 ⋅ (1 − *thr*)^−1^ Hz. Model responses were simulated to a Gaussian white-noise stimulus (60 Hz) for durations between 1 and 20 minutes (in 1-minute steps). For each combination of threshold and stimulus duration, the spatiotemporal RF was re-estimated from the simulated spikes via the spike-triggered average and compared to the ground-truth filter by the root-mean-square error (RMSE) between the estimated and the true spatiotemporal RF, each normalized by its maximum absolute value.

### LNLP model and spike-triggered covariance analysis

To demonstrate that the RF of an ON-OFF cell (see Extended Data Fig. 4 and 5), which cannot be recovered by the spike-triggered average (STA), can be recovered by spike-triggered covariance (STC) analysis, we simulated an ON-OFF model cell and analyzed its responses with STC (Extended Data Fig. 4c,d and Extended Data Fig. 5a,b). The model cell was constructed as the sum of an ON and an OFF Linear-Nonlinear-Poisson subunit. Each subunit had a single-Gaussian spatial filter as defined above, on a 10 × 10 pixel grid, with amplitude *A* = 1, center *µ*_*x*_, *µ*_*y*_) = (5,5, and standard deviations *σ*_*x*_ = *σ*_*y*_ = 1.5 pixels, and a Gabor temporal filter *g*(*τ*) (ON: *p* = 1; OFF: *p* = −1; *c* = 0.65, *l* = 0.4) evaluated at 12 time points, yielding a 10 × 10 × 12 spatiotemporal filter. A rectified linear nonlinearity, *h*(*x*) = m*ax*(0, *x*), and a firing-rate amplitude of *BB* = 100 Hz were used. The ON-OFF response was obtained by adding spike trains of the ON and OFF subunits and computing the instantaneous firing rate of the combined spike train, without any additional nonlinearity, i.e., as a Linear-Nonlinear-Linear-Poisson (LNLP) model. Responses were simulated to a 10 × 10 × 100,000 Gaussian white-noise stimulus; a Gaussian (rather than binary) stimulus was used because STC requires an unbiased WN stimulus (Schwartz et al., 2006). STC analysis was performed following (Schwartz et al., 2006). Briefly, we computed the spike-triggered average (STA) and spike-triggered covariance (STC) of the stimulus segments preceding each spike. The STA was projected out of the STC matrix before computing the eigendecomposition of the STC matrix. The recovered filter was the leading eigenvector, i.e., the eigenvector associated with the largest eigenvalue. The static nonlinearities of the STA and STC filters were estimated by binning the respective generator signal into 22 equally sized bins.

We used STC to analyze RGC responses to the white-noise stimulus. At the full stimulus resolution, the leading and trailing STC eigenvectors were estimated as above and assessed by the RF QI. This procedure recovered essentially no RFs (Extended Data Fig. 5c). We, therefore, reduced the stimulus dimensionality by restricting the analysis to a 3 × 3 pixels spatial window around the RF center and the 10 most recent time lags (90 dimensions; cf. (Ahn et al., 2020)). For cells for which a WN STA with a RF QI > .3 was available, we centered the spatial window for the STC analysis on the WN STA. Otherwise, the spatial window was centered on the RCASE-derived RF center. The significance of the leading and trailing STC eigenvalues was assessed by a permutation test: the spike train was circularly shifted by a random offset, and the STC was recomputed 1,000 times. An eigenvalue was deemed significant if it fell outside the central 99% interval (0.5th–99.5th percentile) of this null distribution. A cell was counted as STC-recovered, when at least one eigenvector was significant and yielded a RF with RF QI > 0.3 (Extended Data Fig. 5d–f); cells with less than one spike per stimulus dimension were excluded.

### Fitting LN-models to RGC responses

We fitted Linear-Nonlinear models (LN-models), consisting of a linear spatiotemporal filter, followed by a static nonlinearity to RGC responses to the WN and RMO stimulus respectively. The models fit to responses to the WN stimulus were used to predict WN responses, while the models fit to responses to the RMO stimulus were used to predict RMO responses. To match the refresh rate of the WN, we downsampled the RCASE stimulus design matrix to 20Hz for mouse experiments and to 30Hz for primate experiments. To determine the RF centers and optimal time lag *τ*_*opt*_ at which the temporal RF was maximal, spatiotemporal RFs were estimated via the spike-triggered average for the WN stimulus and via RCASE for the RMO stimulus. To diminish the effect of overfitting, the dimensionality of the stimulus design matrices *X*_*WN*_ and *X*_*RMO*_ were reduced by cutting an area of 10x10 around the RF center and limiting the number of time lags to 15, resulting in spatiotemporal RFs with a total of 1500 parameters. Time lags to include for the RMO stimulus were set on a cell-by-cell basis as the 7 time bins before and after *τ*_*opt*_, including *τ*_*opt*_ . The dataset was split into a training and test set using 10-fold cross validation (90% training data, 10% test data). In each fold the RFs were estimated by linear regression:

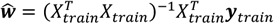

We calculated the model responses for the training and test set as

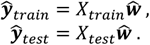

To estimate the static nonlinearity, we followed the approach described in (Chichilnisky, 2001). Briefly, for each cell, we created 20 uniformly sized bins over the range of the generator signal **ŷ**_*train*_. We then averaged spike counts ***y***_*train*_ within these bins, resulting in an empirical estimate for the nonlinearity, to which we fitted a sigmoid,

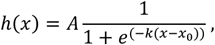

where *A* scaled the sigmoid in the range [0, *A*], *k* controlled the gain of the sigmoid, and *x* was the sigmoid’s midpoint, that is 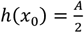. The test error for each fold was computed as the squared Pearson correlation coefficient between measured and modelled response. Reported prediction errors were the average test errors across the 10 folds. Cells with an undefined ρ^2^ in any fold, for example because they did not spike during a test fold, were excluded, leaving n = 1,346 of 1,871 RGCs.

### Clustering

We clustered the light responses of mouse non-direction selective RGCs (n=969, 3 retinae) in response to the RMO stimulus. In these experiments, we parameterized the objects by different sizes, speeds, and colors. For the clustering, each RGC was represented by a feature vector. The feature vector was constructed by concatenating different tuning properties of each cell including the RCASE temporal kernel. Before concatenation to the feature vector, we reduced the dimensionality of the RCASE temporal kernel by scaling it to unit norm, applying PCA, and keeping the PCs associated to the largest eigenvalues which explained 95% of the variance. The feature vector was constructed by concatenating the reduced RCASE temporal kernel (35 features), the RCASE color tuning curve (3 features), the RCASE speed tuning curve (5 features), and the POTH distance tuning curve (20 features). The resulting 63-dimensional feature vector was scaled to unit norm for each cell and arranged into a design matrix, which was used as input to the clustering algorithm. We used Scikit-learn *AgglomerativeClustering* algorithm with linkage *ward* to obtain a hierarchical cluster tree. Here, each neuron started in its own cluster, and pairs of clusters were iteratively merged based on their distance. The number of clusters with more than 30 cells was plotted against the merging step. The algorithm was stopped when the maximum number of clusters with more than 30 cells was reached. Clusters with 30 or fewer cells were discarded. This resulted in 15 clusters (n=795). Population-averaged response features to the RMO, as well as the population- and trial-averaged firing rate to the full-field chirp stimulus (Gaussian Kernel, Δ*t* = 0.01*s*, *σ* = 0.05*s*) were plotted along with the standard error of the mean (SEM) (Extended Data Fig. 6d,e). Clusters were ordered by their average full-field chirp stimulus ON-OFF-index. We visualized the clustering using the nonlinear dimensionality reduction method UMAP (McInnes et al., 2018) (Extended Data Fig. 6c).

---

**Extended Data Fig. 1:**
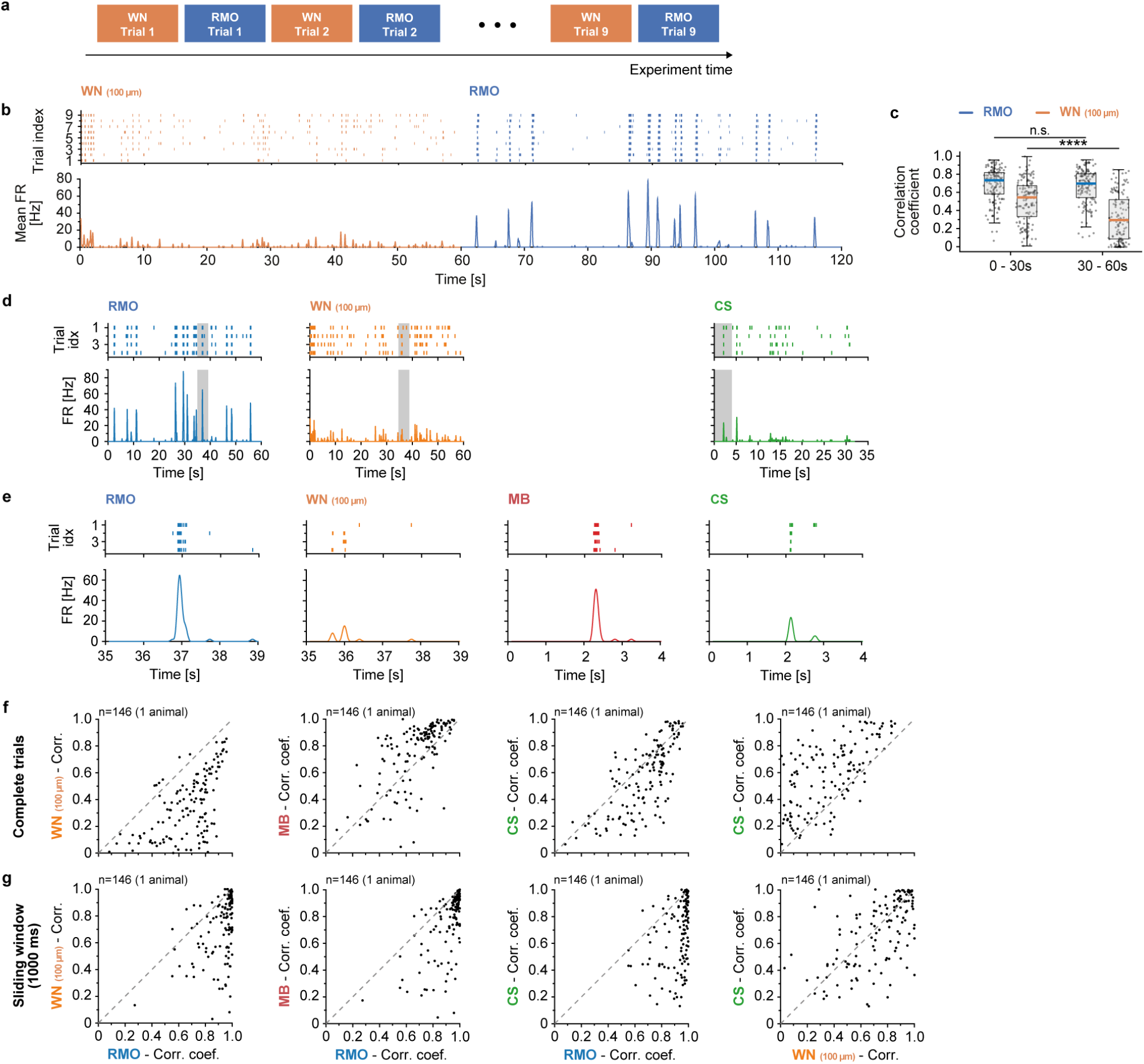
Comparison of response reliability for different stimuli. **a**, Sketch of stimulus protocol. One trial consisted of 1 min WN followed by 1 min RMO. In total there were 9 trials. **b**, Top, Raster plot of an example mouse RGC responding to one trial of WN and one trial of RMO. Bottom, corresponding time-resolved firing rate. **c**, Correlation coefficient computed for the first 30 s and last 30 s for each stimulus (n=146 RGCs, 1 animal). For RMO there was no significant difference (Wilcoxon signed rank test; *P* = 0.0617), whereas for WN the rate dropped (Wilcoxon signed rank test; *P* = 9.7994 . 10^−15^). **d**, Response of an example mouse RGC to the presentation of different stimuli. Top, Raster plot of four repetitions of the same stimulus trial. Bottom, average firing rate over the four trials. Period marked in grey is depicted in e. The moving bar stimulus is not shown, as the whole duration of a trial is shown in **e. e**, Response of an example mouse RGC to the presentation of different stimuli. **f**, Scatter plot of correlation coefficients between the instantaneous firing rates of individual neurons (black dots) across presentations of the same stimulus. The axes represent different stimuli. For a neuron falling below the identity line (dotted line) the correlation between trials for the RMO stimulus was higher than for the comparison stimulus. **g**, Same as f but for a sliding window analysis. Here the correlation coefficient between trials is computed within 1 s windows and then averaged across window positions.

**Extended Data Fig. 2:**
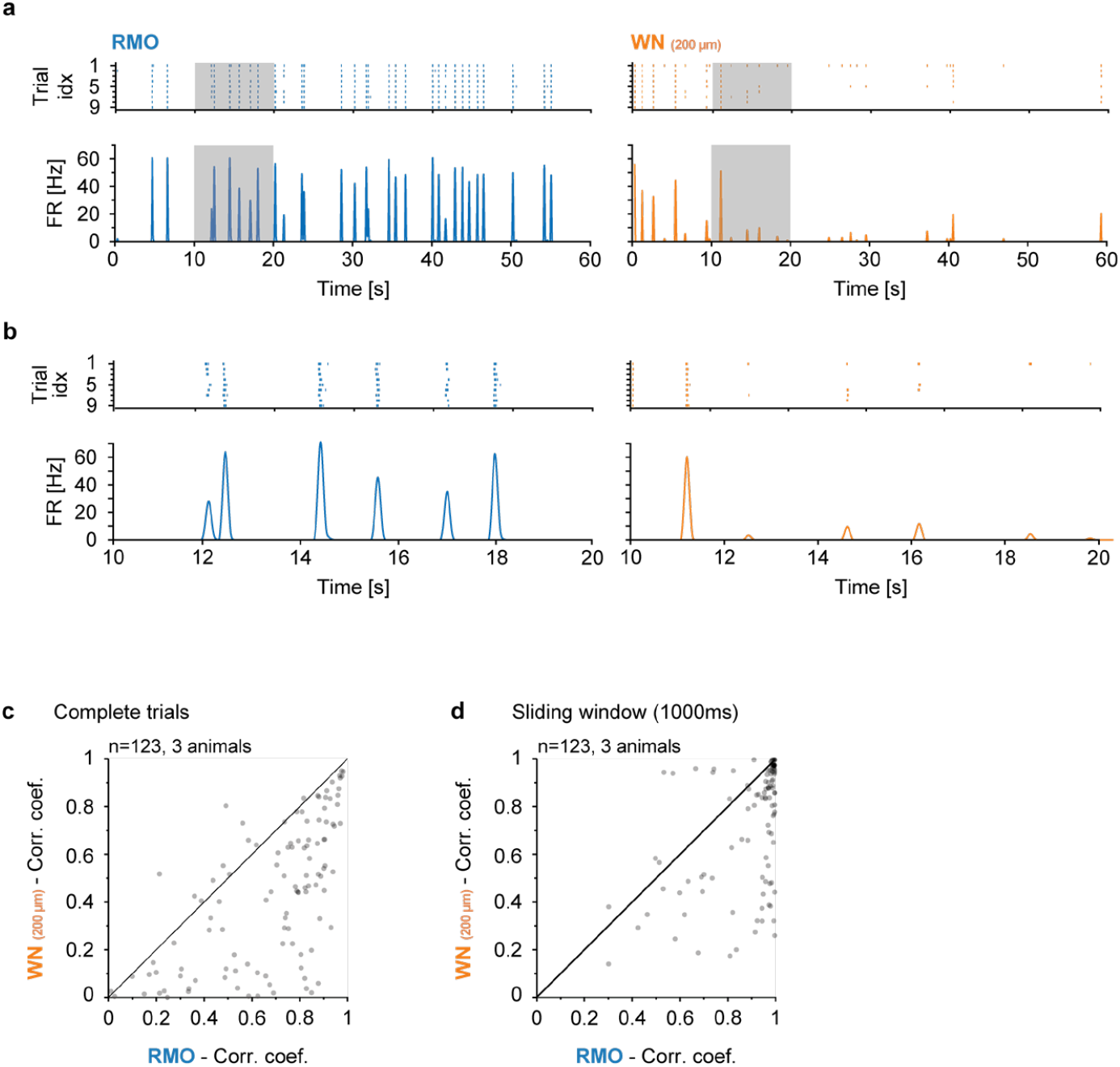
Comparison of spike train reliability between RMO and WN 200. **a**, Response of an example mouse RGC to repeated stimulation with an identical stimulus. Top: Spike trains raster plot (top) and average firing rate (bottom) for nine repetitions (trials) of an identical stimulus presentation of RMO (left) and WN with 200 µm pixel side length (compared to 100 µm pixel side length used in Extended Data Fig. 1). Grey area indicates epoch shown in **b**. **b**, Same as **a** but for a shorter time period. **c**, Scatter plot of correlation coefficients between the instantaneous firing rates of individual neurons (black dots) across presentations of the same stimulus. The axes represent different stimuli. For a neuron falling below the identity line (dotted line) the correlation between trials for the RMO stimulus was higher than for the comparison stimulus. **d**, Same as **c** but for a sliding window analysis. Here, the correlation coefficient between trials is computed within 1 s windows and then averaged across window positions.

**Extended Data Fig. 3:**
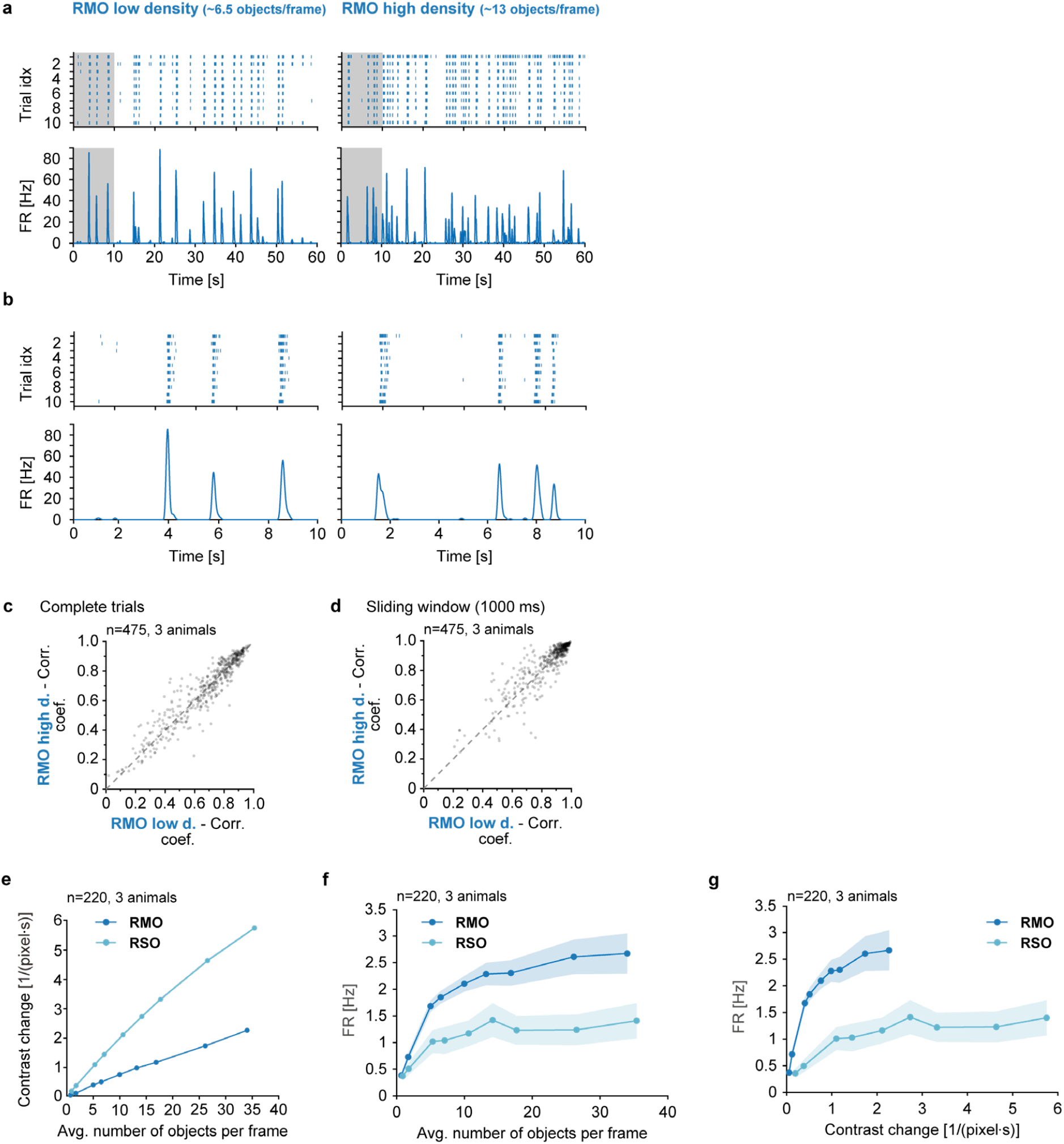
Dependence of RGC responses on object density. **a**, Spiking responses of one example RGC to 10 trials of a low-density and a high-density RMO stimulus (60 s per stimulus). Top: Raster plots of recorded spikes for each trial for the low-density (left) and high-density (right) RMO stimulus. Bottom: Trial-averaged firing rate. **b**, Same as **a**, for the first 10 s of the response (denoted by shadowed area in **a**). **c**, Trial-to-trial response reliability of the recorded RGC population (n=475, 3 mouse retinae), defined as the Pearson correlation coefficient between the binned single-trial firing rates concatenated across all pairs of trials, for the low-density (x-axis) versus the high-density (y-axis) RMO stimulus, computed over the complete trial. Dashed line: identity. **d**, Same as **c**, with the correlation coefficient computed in a 1000-ms sliding window, taking the maximum over window positions. **e**, Contrast change as a function of the average number of objects per frame, for the random moving objects (RMO) and random static objects (RSO) stimuli across nine object densities. Each point corresponds to one object density. **f**, Median firing rate over the recorded RGC population (n=220, 3 mouse retinae) as a function of the average number of objects per frame, for the RMO and RSO stimuli. Shaded area: standard deviation of median firing rates, obtained through bootstrap resampling. **g**, Same as **f**, plotted against the contrast change.

**Extended data Fig. 4:**
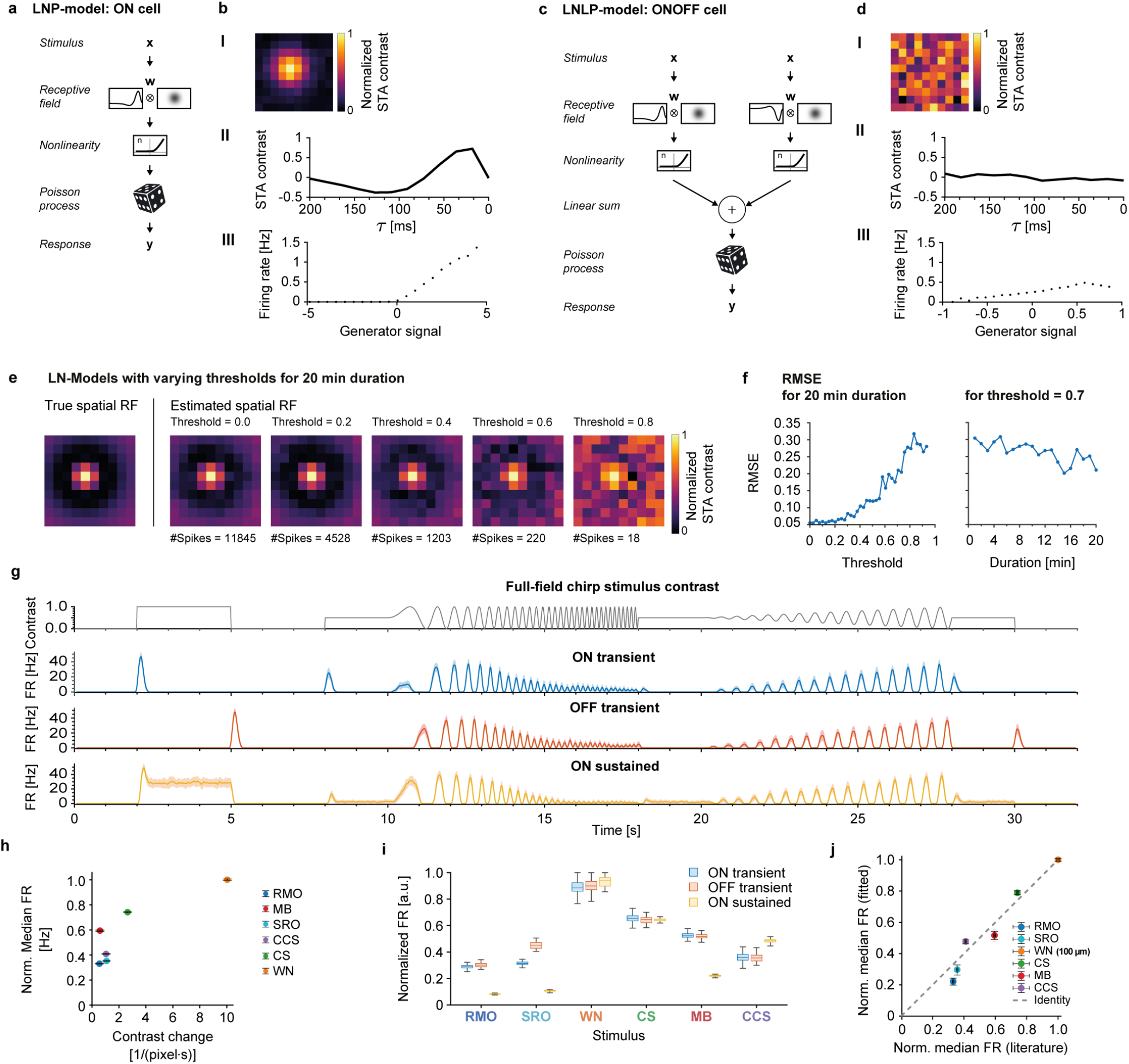
RGC model evaluation. **a**, Schematic of a linear-nonlinear-Poisson cascade model (LNP). The input to the model is a stimulus vector *x*, containing spatiotemporal stimulus intensities. The stimulus is convolved with a spatiotemporal filter (RF). The obtained linear response is passed through a static nonlinearity, and an inhomogeneous Poisson process generates spikes. Spike times are binned in time to obtain the response vector *y*. **b**, STA analysis of the LNP model ON cell depicted in **a** when stimulated with WN (10x10 pixels, 100,000 Frames, ∼28 min at 60 Hz). From top to bottom: I, STA spatial RF; II, STA temporal RF; III, static nonlinearity recovered from the STA. **c-d**, same as a-b but for an LNLP model with an ON- and an OFF-subunit. No STA was recovered from this model. **e**, True and estimated RFs using STA on WN stimulation for an LN model for which the spiking threshold was systematically varied. The higher the spiking threshold, the longer the necessary stimulus duration to estimate the STA. **f**, RMSE of the estimated RF in e for different spiking thresholds (left), and for a fixed threshold but varying durations (right). **g**, Average responses of three different types of LNP-models (n=50) resembling common mouse RGC response behaviors: ON-transient, OFF-transient, and ON-sustained cells. Shaded area is the standard deviation. **h**, Contrast change vs. normalized median firing rates of LNP-model cells (n=110 ON-transient cells). Normalized by maximum of the median firing rates. Error bars: standard deviation of the normalized median firing rates, obtained through bootstrap resampling. **i**, Normalized firing rates of different LNP-model cell types. Normalized within cell type by maximum firing rate (n=110 per cell type). **j**, Normalized median firing rate of LNP-model cells (n=110 ON-transient cells) vs. normalized median firing rate for LNP-models fitted to recorded ON-transient retinal ganglion cells (n=90). Normalized by maximum of the median firing rates within each group. Error bars: standard deviation of the normalized median firing rates, obtained through bootstrap resampling.

**Extended Data Fig. 5:**
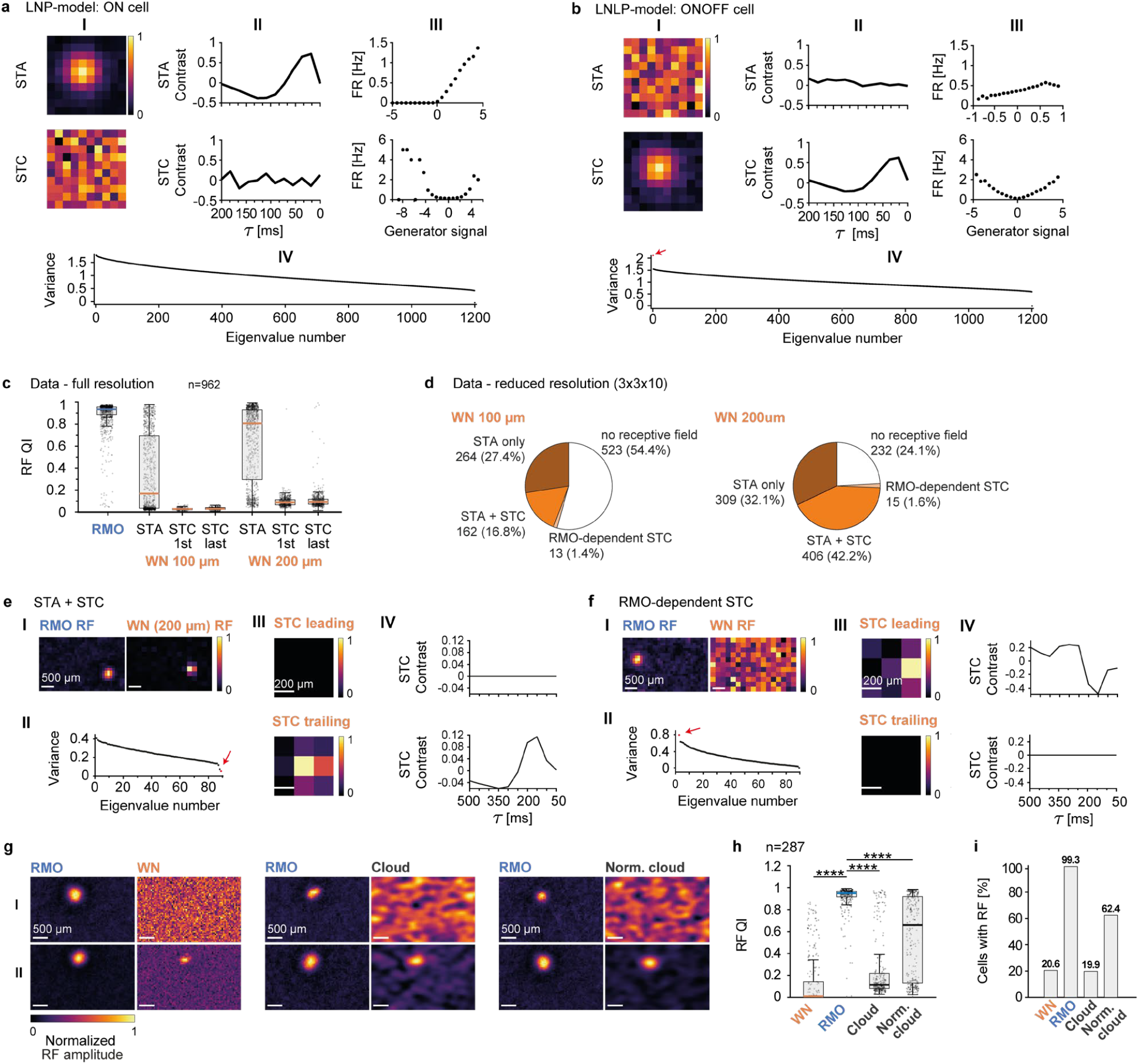
Comparison of different RF mapping approaches. **a**, STC analysis of a linear-nonlinear Poisson (LNP) model ON cell. **I**, Spatial RFs of the spike-triggered average (STA, top) and of the leading STC eigenvector (bottom), each scaled to [0, 1]. **II**, Corresponding temporal RFs. **III**, Corresponding static nonlinearities. **IV**, Sorted eigenvalues of the STC matrix. **b**, Same as **a**, for an LNLP model ON-OFF cell based on the sum of an LNP model ON cell and an LNP model OFF cell. The small red arrow points at the largest eigenvalue. **c**, RF quality index (RF QI) of RFs estimated in the mouse retina with RMO/RCASE and with WN STA and STC analysis (STC; first and last filters), for WN edge size of 100 and 200 µm (n = 962 RGCs, 3 retinae). Boxes show the median and interquartile range with whiskers; individual RGCs are overlaid as jittered points. **d**, Comparison of STA- and STC-identifiable RFs using WN stimulation. Fraction of RGCs assigned to each RF estimation category, for WN of 100 µm (left) and 200 µm (right) edge size (n = 962 RGCs, 3 retinae). Cells are partitioned into four categories: STA only; STA and STC; RMO-dependent STC (an RF recovered by STC but not STA, identifiable only because RCASE located its RF center using the RMO stimulus); and no RF. **e**, STC analysis of an example retinal ganglion cell from the “STA and STC” category (**d**), for a 200 µm WN stimulus. **I**, Spatial RFs estimated with the RMO/RCASE stimulus (left) and with the WN STA (STA, right), each scaled to [0, 1] (scale bar, 500 µm). **II**, Sorted eigenvalues of the STC matrix; the trailing eigenvalue is marked in red. **III**, Spatial RFs of the leading (‘STC leading’ and trailing (‘STC trailing’) STC eigenvectors, each scaled to [0, 1] (scale bar, 200 µm). **IV**, Corresponding temporal RFs of the leading (top) and trailing (bottom) STC eigenvectors. **f**, Same as **e**, for an example cell from the “RMO-dependent STC” category (**d**) whose RF is recovered by STC but not by WN STA. Here the leading STC eigenvector is marked red (**II**); **III** and **IV** show the leading and trailing eigenvector’s spatial and temporal RF. **g**, Spatial RFs estimated with RMO and with three WN stimuli, binary white noise (WN), cloud white noise (cgWN), and normalized-cloud white noise (cngWN), for example mouse retinal ganglion cells. For each white noise stimulus, two example cells are shown: one for which an RF was recovered with RMO (RF QI > 0.3) but not with the corresponding WN stimulus (RF QI < 0.3), and one for which both stimuli yielded an RF (RF QI > 0.3). Within each pair, the RMO RF is shown next to the corresponding RF (WN, cgWN, cngWN), each scaled to [0, 1], scale bar, 500 µm). **h**, RF QI for all recorded RGCs, estimated with RMO/RCASE and the same three WN stimuli as in **g** (WN, cgWN, cngWN) (n = 287 RGCs, 2 retinae). Boxes show the median and interquartile range with whiskers; individual RGCs are overlaid as jittered points. (**** P < 0.0001 for WN vs. RMO, RMO vs. cloud, and RMO vs. normalized-cloud; Wilcoxon signed-rank test, Bonferroni-corrected for three comparisons). **i**, Fraction of recorded RGCs with a recovered RF (RF QI > 0.3), estimated with RMO/RCASE and the same three WN stimuli as in **g** and **h** (WN, cgWN, cngWN) (n = 287 RGCs, 2 retinae).

**Extended Data Fig. 6:**
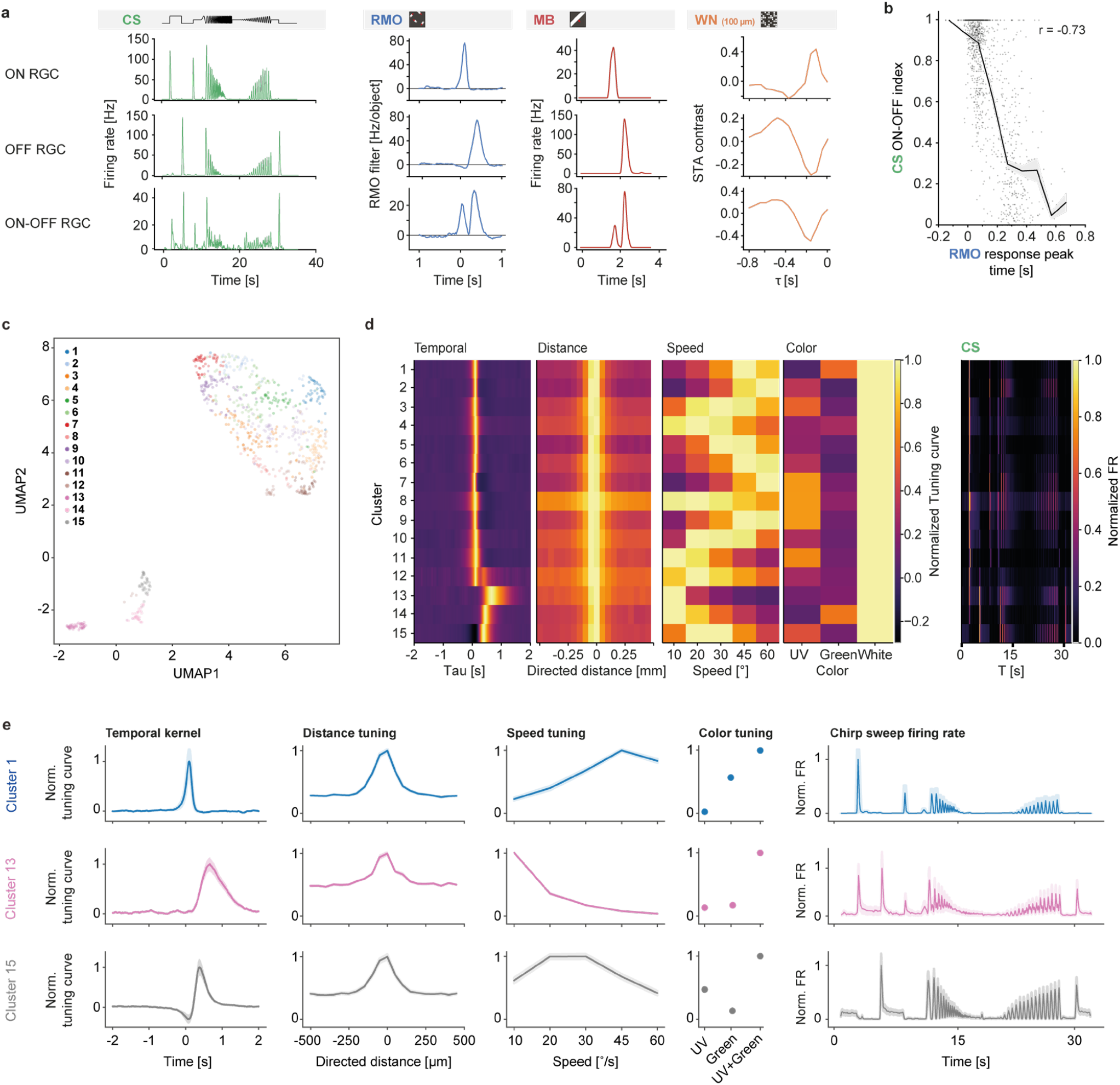
Population response properties estimated with RMO/RCASE. **a**, Average responses of an ON-, an OFF-, and an ON-OFF-RGC to the CS, RMO, MB, and WN stimulus. RMO and WN stimuli: temporal kernel; CS and MB stimuli: trial-averaged firing rates. **b**, Comparison of the ON-OFF index estimated with the CS stimulus and the time of the peak of the RMO temporal kernel. Individual RGCs are depicted as gray dots, the line shows the average CS ON-OFF index within bins (100ms bin size) of the RMO response peak time; shaded area: standard error of the mean. **c**, UMAP projection of mouse RGC response features obtained with RCASE with the RMO stimulus. **d**, Cluster-averaged neural response features obtained with RCASE and the RMO stimulus, including temporal kernel, distance tuning curve, speed tuning curve, and color tuning curve, for 15 clusters obtained via hierarchical clustering (Mouse retina, n=795, 3 retinae). **e**, Clustering using RMO response features (leftmost 4 columns): temporal kernel, distance tuning curve, speed tuning curve, and color tuning curve. Shown are normalized average response features of three example clusters out of a total of 15 as well as the normalized trial- and population-averaged firing rate in response to the chirp sweep (not used for clustering); shaded area: standard error of the mean.

**Extended Data Fig. 7:**
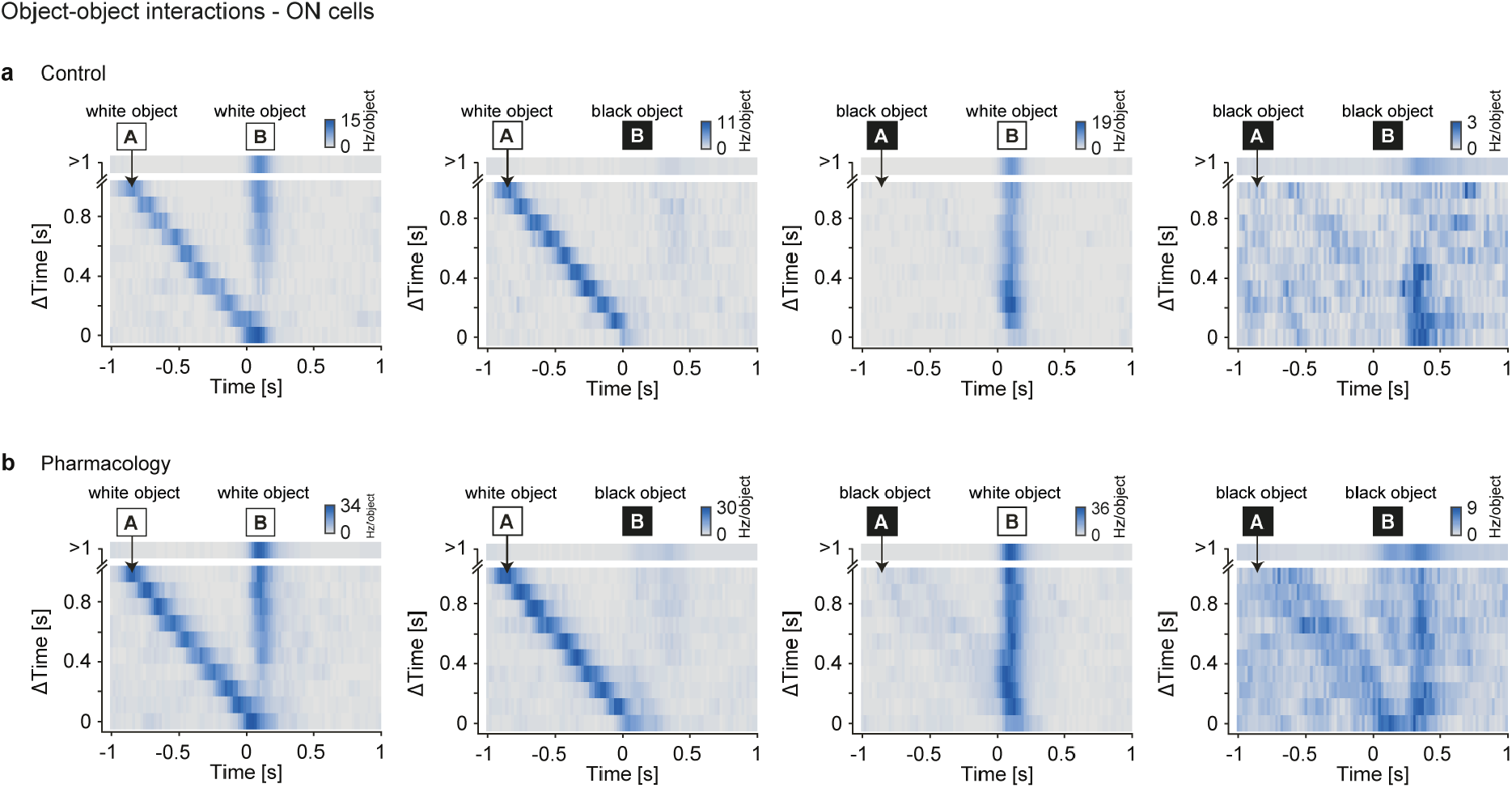
Response analysis to pairs of objects. 2D-Histograms (t, Δt) of responses towards pairs of objects in different conditions (black vs. white objects, control vs. pharmacological block of amacrine-cell mediated inhibition (strychnine, gabazine, picrotoxin)). Object A arrives first, and responses are aligned with respect to the time when object B is closest to the RF center. Δt is the time difference between the times of closest approach to the RF center for object A and object B. t is the time of the response with respect to the closest approach time point for object B. **a**, Control condition; **b**, pharmacological block of inhibition. Columns show 4 different combinations of black and white object pairs.

**Extended Data Fig. 8:**
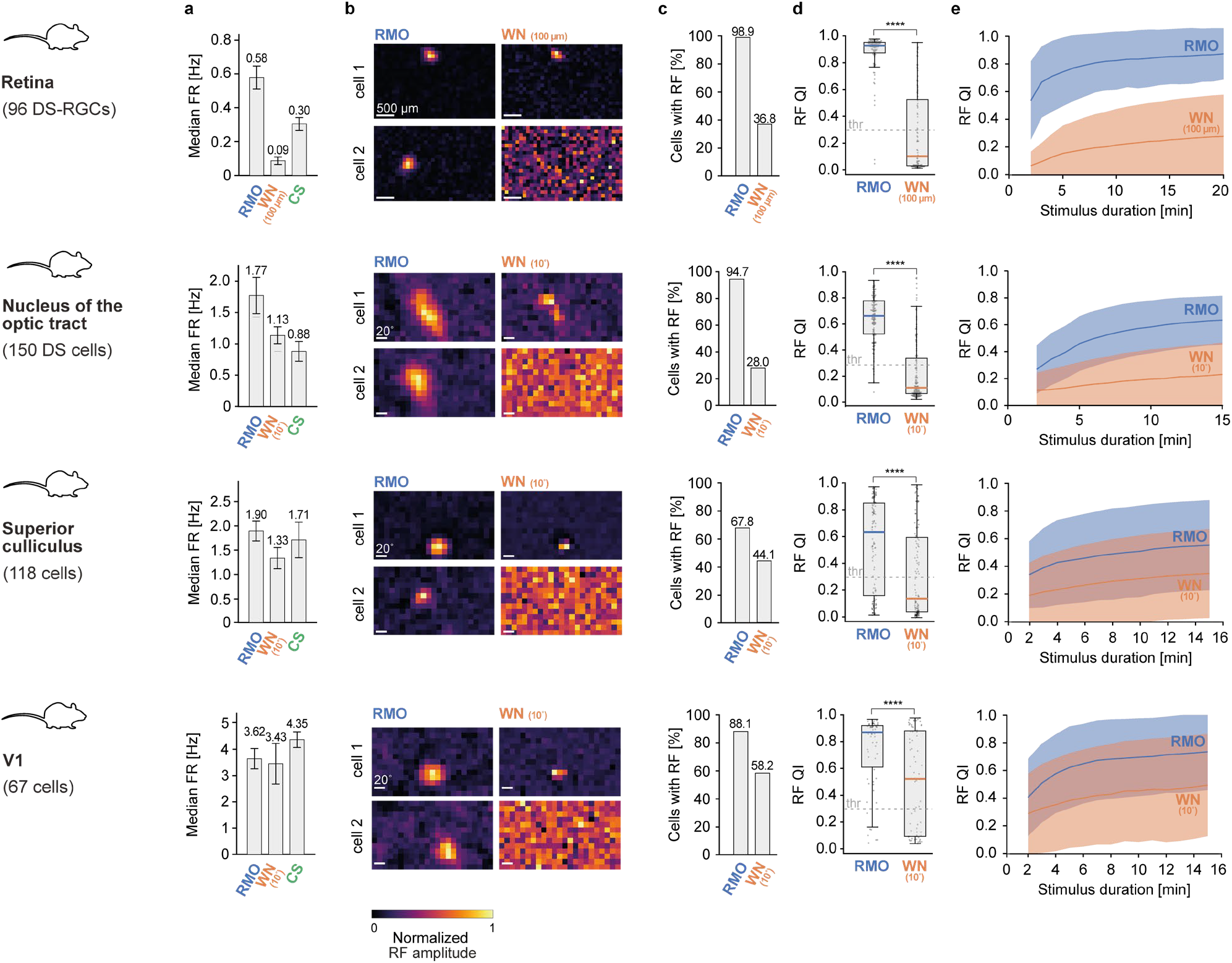
Comparison of RMO & WN RF mapping across processing stages in the mouse retina & cortex. **a-e**, from top to bottom: Mouse DS-RGCs (n=96, 4 retinae); DS-Cells in NOT (n=150, 7 mice); cells in the SC (n=118, 3 mice); cells in V1 (n=67 cells, 4 mice). **a**, Median firing rate for different light stimuli. Error bars: standard deviation of median firing rates, obtained through bootstrap resampling. **b**, Spatial RFs for RMO and WN for two example cells. Same panels as in Fig.6a-d II. **c**, Fraction of cells with RF quality index *RF*_*QI*_ > 0.3 for RMO and WN. **d**, *RF*_*QI*_ for the population of recorded RGCs *(**** P < 0*.*0001, Wilcoxon signed-rank test)*. **e**, *RF*_*QI*_ as a function of stimulus duration.

**Extended Data Fig. 9:**
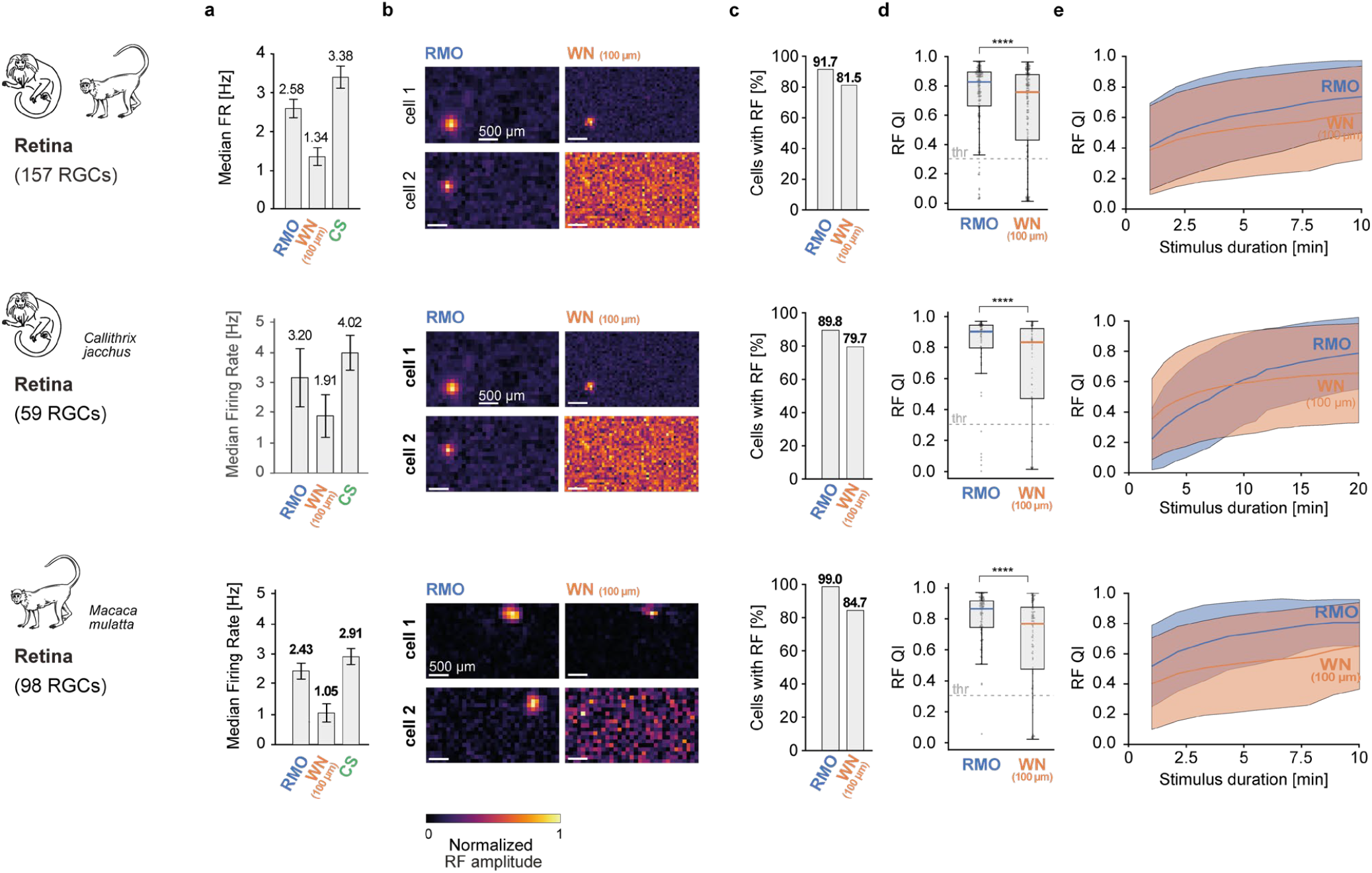
Comparison of RMO & WN RF mapping in the primate retina separated by species. **a-e**, from top to bottom: all primate retinas combined (n=157, 2 retinas); Marmoset retina only (n=59, 1 retina); Macaque retina only (n=98, 1 retina). **a**, Median firing rate for different light stimuli. Error bars: standard deviation of median firing rates, obtained through bootstrap resampling. **b**, Spatial RFs for RMO and WN for two example cells. **c**, Fraction of RGCs with RF quality index *RF*_*QI*_ > 0.3 for RMO and WN. **d**, *RF*_*QI*_ for the population of recorded RGCs *(**** P < 0*.*0001, Wilcoxon signed-rank test)*. **e**, *RF*_*QI*_ as a function of stimulus duration.

**Extended Data Fig. 10:**
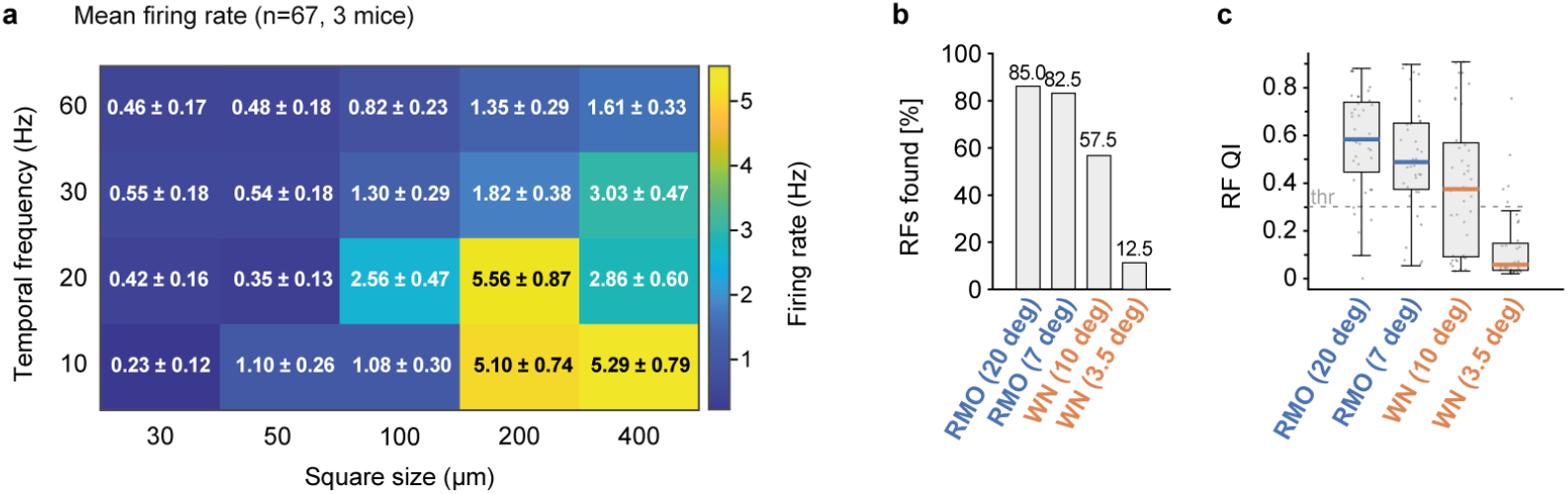
Dependence of the quality of RF estimation on stimulus parameters and stimulus choice in the retina and NOT. **a**, Mean firing rate of mouse retinal ganglion cells in response to a white noise (WN) stimulus, as a function of checkerboard square size and temporal frequency. Color encodes the population mean firing rate; each cell is annotated with the mean ± standard error of the mean (n=67, 3 mice). **b**, Fraction of NOT neurons with a RF quality index *RF*_*QI*_ > 0.3, shown for RMO and WN stimuli at two object sizes each (n = 40 cells, 4 mice). **c**, Distribution of *RF*_*QI*_ across the same neurons as in **b** for each stimulus condition; boxes show the median and interquartile range with whiskers, and overlaid points show individual cells.

**Extended Data Table 1:** Mouse RGC types and preferred light stimulus size, centered on their RF. The maximum light stimulus size triggering a response of at least 50% of the maximum signal and expected response to a 1 mm^2^ full contrast light flash. Intrinsically photosensitive RGCs are not included in the list.

| Mouse RGC cell type |  | Spot size of the best response [μm] | Spot size with 50% response [μm, estimated] | RGC responds to uniform stimuli covering 1 mm<sup>2</sup> or more | Reference |
| --- | --- | --- | --- | --- | --- |
| Cell types responding to large uniform light stimuli | tON α | ~ 500 | 700 | Yes | (Krieger et al., 2017; Farrow et al., 2013) |
|  | sON α | 350 | 1200 | Yes | (Estevez et al., 2012; Krieger et al., 2017) |
|  | tOFF α | ~ 600 | 1000+ | Yes (assumed) | (Farrow et al., 2013; Warwick et al., 2018) |
|  | sOFF α | ~ 250 | 1200 | Yes | (Farrow et al., 2013) |
|  | SbC | 200 | 600 + | Yes (assumed) | (Tien et al., 2015) |
|  | PixON | 300 | 1200 | Yes | (Johnson et al., 2018) |
|  | vOS ON large | 220 | 800 | Yes | (Nath & Schwartz, 2016) |
|  | hOS ON large | 220 | 800 | Yes | (Nath & Schwartz, 2016) |
|  | OFF vOS | 220 | 800 | Yes | (Nath & Schwartz, 2016) |
|  | oDS (3) | 300 | 700 | Yes | (Dhande et al., 2013) |
| Cell types not responding to large uniform light stimuli | F-mini ON | 60 | 160 | No (assumed) | (Rousso et al., 2016) |
|  | F-mini OFF | 75 | 160 | No (assumed) | (Rousso et al., 2016) |
|  | F-midi ON | 90 | 160 | No (assumed) | (Rousso et al., 2016) |
|  | F-midi OFF | 100 | 130 | No | (Rousso et al., 2016) |
|  | w3 | 120 | 190 | No | (Farrow et al., 2013; Kim et al., 2010; Zhang et al., 2012) |
|  | JamB | 125 | 250 | No | (Farrow et al., 2013; Joesch & Meister, 2016) |
|  | HD1 | 110 | 240 | No | (Jacoby & Schwartz, 2017) |
|  | HD2 | 130 | 200 | No | (Jacoby & Schwartz, 2017) |
|  | UHD | 90 | 180 | No | (Jacoby & Schwartz, 2017) |
|  | ooDS (4) | 210 | 350 | No | (Dhande et al., 2013) |
|  | vOS ON small | ~180 | ~300 | No | (Nath & Schwartz, 2016) |
|  | hOS ON small | ~180 | ~300 | No | (Nath & Schwartz, 2016) |

**Extended Data Table 2:** Properties of the different RMO stimuli used in this study. Object size, speed, density (mean number of objects per frame), object and background colors, and stimulus duration are listed for each experimental condition. Where several values are listed, the corresponding conditions were presented within the same experimental condition.

|  | Object size [um] | Object speed [um/s] | Object density [#objects/frame] | Object colors | Background color | Duration [min] |
| --- | --- | --- | --- | --- | --- | --- |
| Mouse retina<br>WN comparison | 200 | 1296 | 7.1 | White | Black | 20 |
| Mouse retina<br>Clustering | 75, 100, 125, 150, 200 | 288, 576, 864, 1296, 1728 | 26 | UV, Green, White | Black | 90 |
| Mouse retina<br>Object-object interaction | 200 | 1296 | 9.1 | White/Black | Gray | 40 |
| Mouse retina<br>Response Reliability | 100, 200 | 1296 | 6.5 | White | Black | 1 |
| Mouse retina<br>Stimulus density WN comparison | 200 | 1296 | 6.5, 12.5 | White | Black | 1 |
| Mouse retina<br>Stimulus density comparison | 200 | 1296 | 0.8, 1.7, 5.2, 6.8, 10.3, 13.7, 17.3, 26.3, 34.7 | White | Black | 2 |
| Marmoset retina | 150, 200, 250, 300 | 192, 479, 959, 1918, 2877, 4315 | 24.4 | White | Black | 20 |
| Macaque retina | 200 | 1296 | 20.7 | White | Black | 10 |
| Nucleus of the optic tract (NOT) | 576 | 1296 | 5.6 | White | Black | 15 |
| Primary visual cortex (V1) | 576 | 1296 | 5.6 | White | Black | 15 |
| Superior colliculus (SC) | 576 | 1296 | 5.6 | White | Black | 15 |

